# Decoupling tRNA promoter and processing activities enables specific Pol-II Cas9 guide RNA expression

**DOI:** 10.1101/342485

**Authors:** David JHF Knapp, Yale S Michaels, Max Jamilly, Quentin RV Ferry, Hector Barbosa, Thomas A Milne, Tudor A Fulga

## Abstract

Spatial/temporal control of Cas9 guide RNA expression could considerably expand the utility of CRISPR-based technologies. Current approaches based on tRNA processing offer a promising strategy but suffer from high background. Here we developed a variant screening platform to identify differential sequence determinants of human tRNA promoter and processing activities. Rational design based on the ensuing principles allowed us to engineer an improved tRNA scaffold that enabled highly specific guide RNA production from a Pol-II promoter.

---

Most CRISPR/Cas9 guide RNA (gRNA) expression systems use RNA polymerase-III (Pol)-III promoters such as U6^1,2^. While highly efficient, these promoters act in a constitutive fashion^3^. An optimized system for tissue specific or inducible gRNA expression could greatly increase the flexibility of the CRISPR/Cas9 system. However, most expression systems enabling spatial/temporal control of promoter activity require Pol-II mediated transcription. Following transcription, Pol-II products are rapidly modified with a 5’ cap and poly-A tail and exported from the nucleus. These modifications and altered localization could prevent efficient use of Cas9 gRNAs^4^. Consequently, a number of strategies have been proposed to excise gRNAs from Pol-II transcripts. These include the use of alternative transcriptional terminators^5^, embedding the gRNA in a spliced intron^6^, self-cleaving ribozymes-based release systems^4,7^, and the use of Csy4 or orthologous ribonucleases^4^. These strategies however, suffer from relatively poor activation rates downstream of Pol-II promoters, or require the addition of toxic^4^ and potentially immunogenic proteins. Thus, there remains a need for the development of effective non-constitutive CRISPR/Cas9 gRNA expression systems.

tRNAs represent a highly conserved class of RNA molecules which are recognized and precisely processed by RNase P and RNase Z^8^. Various tRNAs have been exploited to allow polycistronic gRNA production with high processing efficiencies^9,10^. tRNAs however, contain internal Pol-III promoters^8^. Indeed, tRNAs have been used to replace U6 promoters for gRNA production^11^, albeit at somewhat lower efficiency^12^. Intriguingly, previous studies reported Pol-II specific gRNA activity using tRNA-based multiplexing systems^7,13^. However, in one instance the Cas9 was also placed under inducible control^13^ and thus the gRNA may still have been constitutively expressed. The second study employed two gRNAs flanked by ribozymes, and the detection system relied on releasing both gRNAs^7^. In this case, the first gRNA was upstream of the tRNA and thus not constitutively expressed by the tRNA Pol-III promoter activity. While such a system could potentially allow Pol-II specificity in some cases, this approach would be difficult to generalize.

The regions involved in tRNA promoter and processing activity have been previously identified^14–16^. While most positions overlap, we hypothesized that the differential requirements for these processes (DNA sequence identity and RNA structure for promoter and processing, respectively) might enable their decoupling, and thus provide an opportunity to re-engineer a tRNA scaffold with optimal parameters for gRNA release. To test this hypothesis, we performed a mutational screen on the human tRNA^Pro^ (AGG; tRNAscan-SE ID: chr1.trna58) and independently measured the effects of base substitutions on promoter, as well as 5’ and 3’ processing activities. Based on this screen, we engineered new tRNA variants which have no detectable promoter activity but retain sufficient processing to allow specific Pol-II dependent Cas9 gRNA production.

First, we investigated the transcriptional activity, 3’ processing ability, and functional gRNA production in human cells of several wild-type tRNAs which have been previously used for gRNA multiplexing or Pol-II expression^7,9,11,13^ (Supplementary Fig. 1). This revealed functionally equivalent gRNA generation between the U6 and the tested tRNAs, only falling off slightly with rice tRNA in human cells (Supplementary Figures 1, 2, 3a, Online Methods). These results confirm that tRNAs alone produce functional gRNAs constitutively and independent of external promoters, making them unsuitable for generalizable spatial/temporal controlled expression.

To test whether the processing and promoter activities of human tRNAs could be dissociated, we designed a variant screening strategy using the human tRNA^Pro^ backbone. This entailed generation of high-content libraries in which each construct represented a single variant tRNA^Pro^ flanked by a pair of gRNAs (Fig. 1a, Supplementary Fig. 4). The regions subjected to mutations were chosen based on their involvement in promoter and processing activities, as well as their lack of secondary structure determinants^14–16^. Two parallel libraries were generated, of which one contained an upstream Pol-II CMV promoter and one did not ((+)CMV or (-)CMV). Using an RNA circularization-nested RT-PCR protocol (Supplementary Fig. 1d, 2, see Online Methods) we then sequenced both the pDNA library and the circular RNA (circRNA) products (Fig. 1a). Quantitative analysis of barcode reads in the processed and unprocessed fractions provided an estimate of processing activity, while comparing the abundance of each mutation in the plasmid pool allowed an estimate of promoter strength (Fig. 1a).

**Figure 1.**
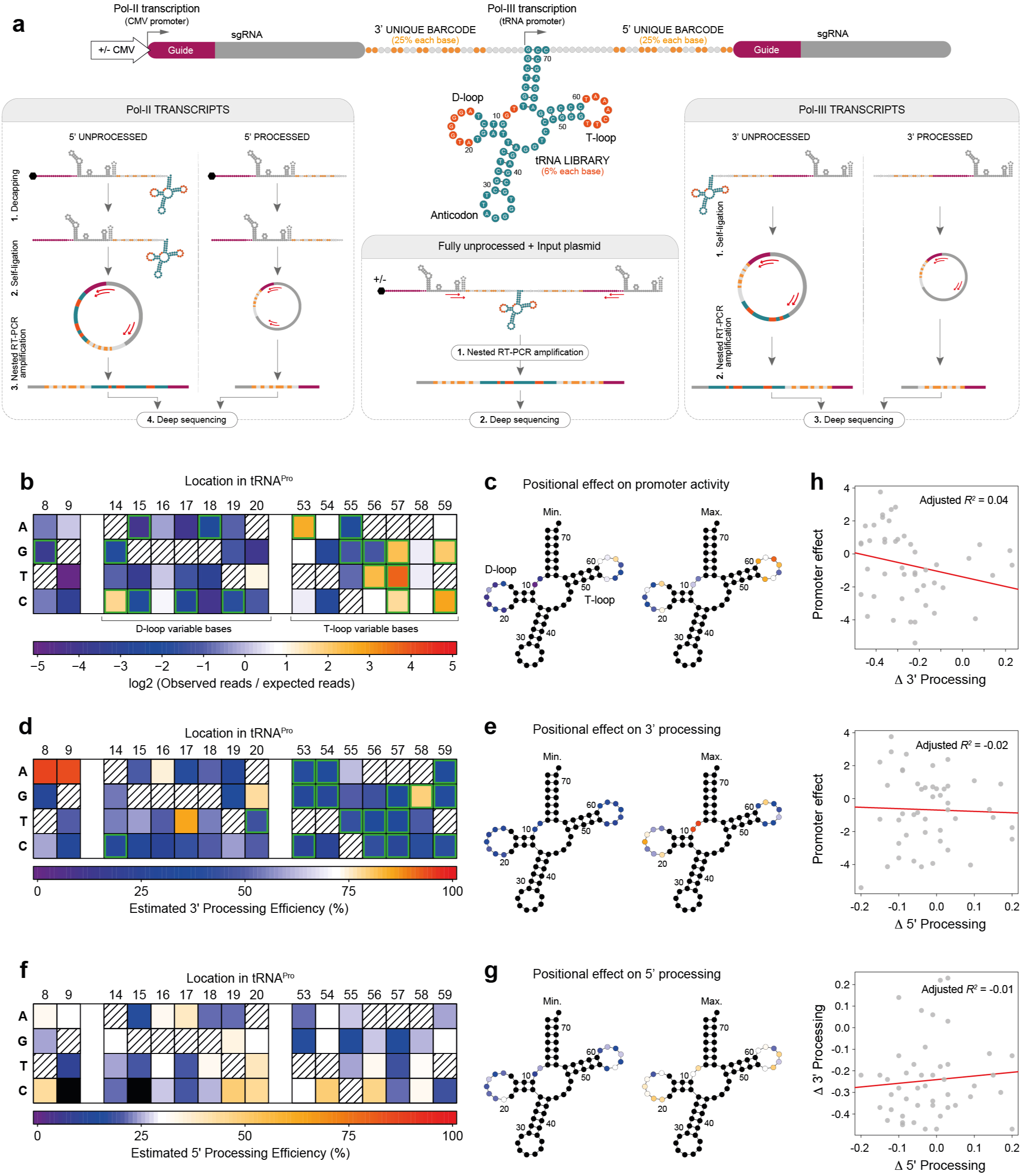
Mutagenesis screen identifies sequence determinants of tRNA processing and promoter activities. **(a)** Experimental design for a tRNA mutagenesis screen in human cells. Parallel libraries of partially degenerate tRNA^Pro^ were created (dark orange = mutated positions), with and without a CMV promoter. Degenerate barcodes (light orange) were placed between the gRNA and tRNA sequences to allow variant identification following processing. A short buffer sequence was included between the barcode and the tRNA to protect the barcode from cleavage. RNA species from (+)CMV libraries were decapped to avoid 5’cap-mediated inhibition of RNA circularization. **(b, d, f)** Heatmaps showing the effects of all single mutations at each nucleotide position on promoter activity (n = 3 paired pDNA/circRNA libraries with no Pol-II promoter) (**b**), 3’ processing (n = 3 paired pDNA/circRNA libraries with no Pol-II promoter) (**d**), and 5’ processing (n = 3 paired pDNA/circRNA libraries with CMV promoters) (**f**) (hatched squares = wild-type nucleotides; black squares = no measurement). In **(b)** promoter activity was calculated as the ratio of observed reads in the RNA fraction of a given mutation compared to its expected frequency in the library from sequencing the plasmid DNA (green borders = changes from wild-type with a probability of 80% or greater; Bayesian Estimation Supersedes the t-test “BEST” test^20,21^; only mutations with observations in all 3 libraries were included in significance testing). In **(d, f)** processing efficiency was inferred by averaging the binomial probability distributions (processed of total reads) across replicates then taking the point of highest probability as the final value estimate (green borders = changes from wild-type with probability densities overlapping by less than 5%). **(c, e, g)** Corresponding tRNA diagrams showing for each modified position, the mutation which rendered the lowest (left) and highest (right) levels of their respective measurements (colors correspond to the heatmaps in b, d, f, respectively). **(h)** Correlation plots for all combinations of measurements. Specific nucleotide changes are as indicated. Red lines reflect a linear fit.

Analysis of promoter strength revealed that most promoter inactivating mutations resided in the D-loop, although position- and even nucleotide-specific effects were observed across all variable sites (Fig. 1b, c). In contrast, only a few specific mutations in the T-loop were detrimental to promoter activity, while others seemed to increase it (Fig. 1b, c). With regard to 3’ processing, most (but not all) mutations in the T-loop had strong detrimental effects on processing, while mutations in the D-loop appeared to have a lesser impact (Fig. 1d, e). Estimations of 5’ processing were hampered by barcode degradation in these libraries, presumably due to the decapping reaction. Therefore, only a partial barcode could be recovered, limiting the number of reads available for analysis and artificially lowering processing estimates. Nevertheless, values were obtained for most single nucleotide variants. This identified several specific nucleotides both in the D- and T-loops which appear to affect 5’ processing (Fig. 1f, g). Interestingly, there was little correlation between mutations affecting 3’ processing, 5’ processing, and promoter activity, supporting our hypothesis that these activities could be dissociated to some degree (Fig. 1h).

Based on the results of our mutagenesis screen, we selected pairs of mutations which should maximally decrease promoter strength while minimally affecting processing ability, as well as pairs which should inhibit processing but not affect promoter strength (Fig. 2a). Since most promoter-detrimental mutations mapped to the D-loop and these tended to have lesser effects on processing, we also created a minimal tRNA backbone by completely deleting the D-loop and the anticodon (ΔtRNA^Pro^, Fig. 2b). This architecture is supported by previous reports suggesting that a similar minimal scaffold retains processing activities equivalent to wild type tRNAs in *Drosophila*^15^. Analysis of the 3’ processing efficiency revealed that all selected double mutants lost their activity to some degree (Supplementary Fig. 5a, b). Surprisingly, in the ΔtRNA^Pro^ scaffolds, which retained enough promoter activity to be detectable, 3’ processing was not decreased compared to wild-type (Supplementary Fig. 5a, b).

**Figure 2.**
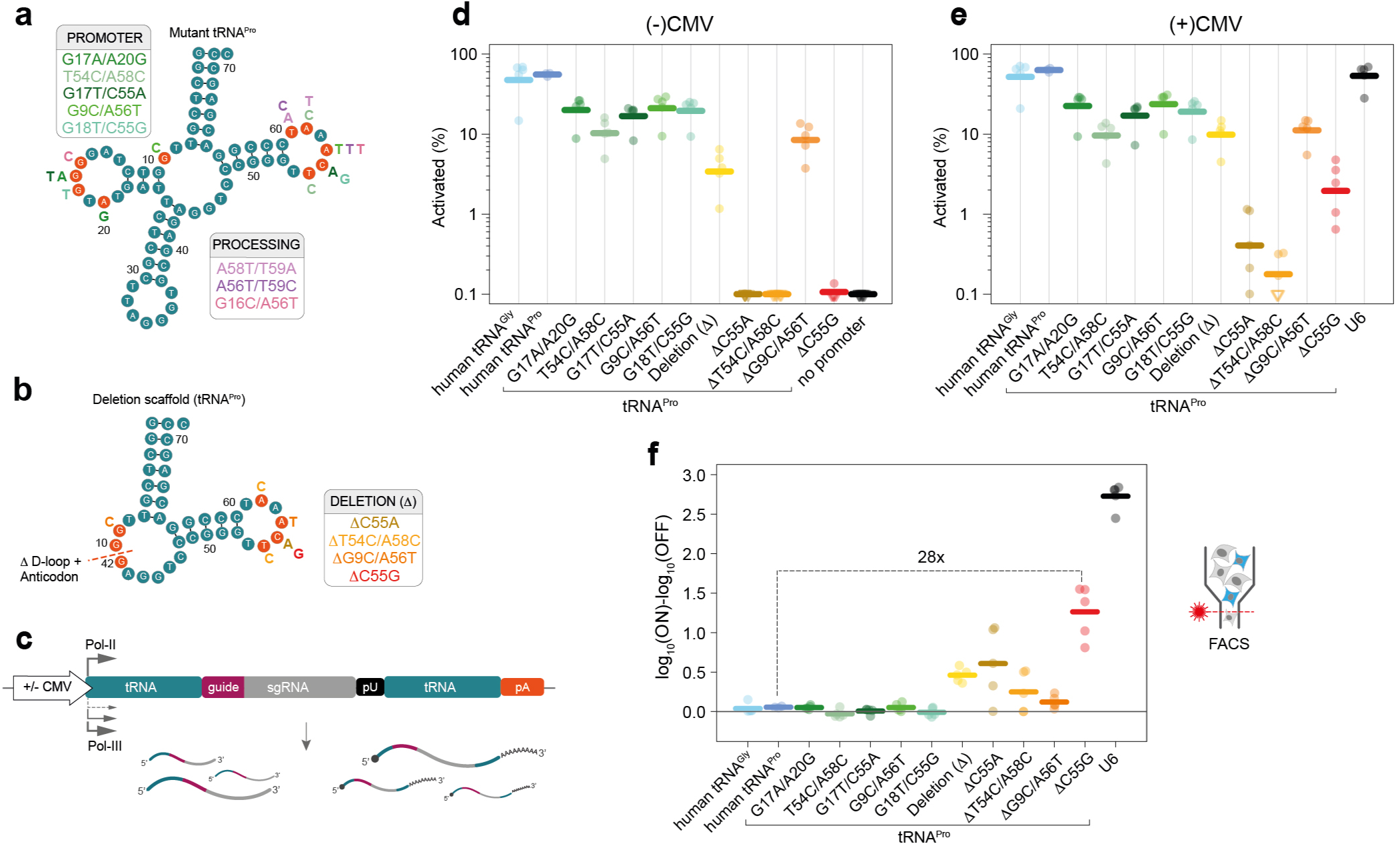
Combinatorial tRNA deletions and point mutations improve the ON-OFF specificity of Pol-II gRNA expression. **(a, b)** tRNA diagrams with mutation pairs predicted from sequencing data to affect 3’ processing only (pink tones), or promoter activity with minimal effect on processing (green/orange/red tones) (left panel). Selected pairs of mutations are shown as letters next to the wild-type **(a)** and ΔtRNA^Pro^ **(b)** scaffolds. **(c)** Constructs used to test ON/OFF ratios of each variant combination in the presence and absence of a Pol-II promoter. **(d, e)** Percentage reporter ECFP^+^ cells within transfected cells with **(d)** and without **(e)** a CMV promoter. Thick lines indicate geometric mean values. Each point represents an independent experiment (n = 4-5). **(f)** Log_10_(% ECFP^+^ cells) in the ON condition compared to OFF condition for each tRNA and U6 (relative to no promoter) (thick lines = mean values; n = 3-5 independent experiments). The U6 and no promoter control are shared with Supplementary Fig. 1e, f and 5d. ΔC55G had a >99.9% probability of decreased activity compared to U6, but a 99% probability of increase compared to parental tRNA (paired BEST tests).

Measurement of promoter activity by qPCR revealed a slight effect of the double mutants designed to decrease promoter strength compared to wild-type (green tones in Supplementary Fig. 5c). Consistently, mutations designed to only affect processing, did not affect promoter activity (pink tones in Supplementary Fig. 5c). The ΔtRNA^Pro^ showed a very strong decrease in promoter activity, and this effect was further exanced in the ΔC55A, ΔT54C/A58C and ΔC55G variants (Supplementary Fig. 5c). Functional gRNA assays revealed a minor decrease in activity for double mutants affecting processing, consistent with their decreased 3’ processing activity (Supplementary Fig. 5d). Double-mutations designed to reduce promoter activity showed an intermediate decrease in functional gRNA activity, consistent with the combination of their decreased promoter strength and partial loss of processing (Supplementary Fig. 5d). The ΔtRNA^Pro^ scaffold showed a strong loss of background gRNA activity, and the addition of other candidate mutations from our screen completely abrogated this leakiness in 3 out of 4 tested combinations (ΔC55A, ΔT54C/A58C and ΔC55G, Supplementary Fig. 5d). These results validate the predictions of our screen, but also suggest that additional synergistic effects may be possible by combining multiple mutations.

Having identified a number of tRNA variants that are potentially competent for gRNA excision from Pol-II transcripts and display reduced or no background activity, we next created paired constructs whereby gRNAs were flanked by engineered tRNAs (Fig. 2c, Supplementary Fig. 6a). We then measured the aggregate processing ability by circularization in the presence a CMV promoter (ON state). This revealed only a slight decrease in overall 3’ + 5’ processing compared to 3’ processing alone, in both wild type tRNA and double mutants, suggesting that the 5’ processing activity was not substantially impaired (Supplementary Fig. 2, 6b). In contrast, the ΔtRNA^Pro^ scaffold showed significantly decreased processing in this assay suggesting that 5’ processing is impaired when the D-loop is entirely removed. Introducing other selected mutations further decreased processing, as expected from their effects on 3’ processing. Several combinations however, retained a readily detectable degree of processing. In particular, the ΔC55G tRNA^Pro^ displayed the highest processing ability (Supplementary Fig. 6b) amongst combinations devoid of leakiness (Fig. 2d and Supplementary Fig. 5d). Quantification of gRNA levels showed strong ON-OFF ratios in all ΔtRNA^Pro^ scaffold combinations (Supplementary Fig. 6c). Interestingly, RNA levels did not increase when a Pol-II promoter was added in front of the wild-type tRNA (Supplementary Fig. 6c) consistent with reports that active Pol-III promoters may inhibit nearby Pol-II activity^17,18^.

Next, we tested the levels of functional gRNAs in each case and determined the reporter activation in the ON and OFF states. This analysis revealed poor ON-OFF ratios for wild-type tRNAs as well as the double mutants (Fig. 2d-f), as predicted by their high background expression and negligible increase in RNA abundance in the presence of a Pol-II promoter. In contrast, the ΔtRNA^Pro^ scaffold and derivatives showed substantially improved ON-OFF ratios due to decreased or absent background activation (Fig. 2d). Importantly and consistent with our promoter and processing assays, while still lower than U6, the ΔC55G tRNA^Pro^ had an ON-OFF ratio over an order of magnitude higher than the wild type tRNA^Pro^ (Fig. 2f).

To establish whether our tRNA deletion/mutant framework is generalizable, we introduced our top performing ΔC55G modification in a human tRNA^Gly^ backbone (GCC; tRNAscan-SE ID: chr1.trna34). This analysis revealed similar elimination of background activity, and improved ON-OFF ratios as observed with the ΔC55G tRNA^Pro^ (Supplementary Fig. 7). These results suggest that the principles described here can be applied to other tRNAs, which in combination could decrease the risk of recombination for multiplexed gRNA frameworks.

Finally, we sought to benchmark our engineered tRNA scaffold against other systems previously employed for Pol-II transcribed gRNA excision^5–7,19^. As reported, all these systems were devoid of significant background activity (Supplementary Fig. 8a). However, in our hands, most of these platforms displayed minimal gRNA-mediated transcriptional activation, except for Csy4 which showed only slightly lower potency compared to our top performing ΔC55G tRNA^Pro^ scaffold (Supplementary Fig. 8b, c).

In this study we identified the base dependencies of human tRNAs promoter activity, 3’ processing and 5’ processing, and used this information to rationally engineer a tRNA scaffold with substantially improved specificity for Cas9 gRNA expression from Pol-II promoters. This framework overcomes the limitations of previous tRNA-mediated release systems, which were compatible with multiplex gRNA delivery but not with spatial/temporal control of gRNA expression, due to their intrinsic Pol-III promoter activity. Further studies will be required to better understand the effects of multiple mutations/deletions in human tRNAs, which might improve the processing activity without loss of specificity. These findings provide new insights into the functional characteristics of human tRNAs and advance existing tools for inducible Cas9 gRNA expression, thus enabling the implementation of more complex research applications.

## FIGURE LEGENDS

**Supplementary Figure 1.**
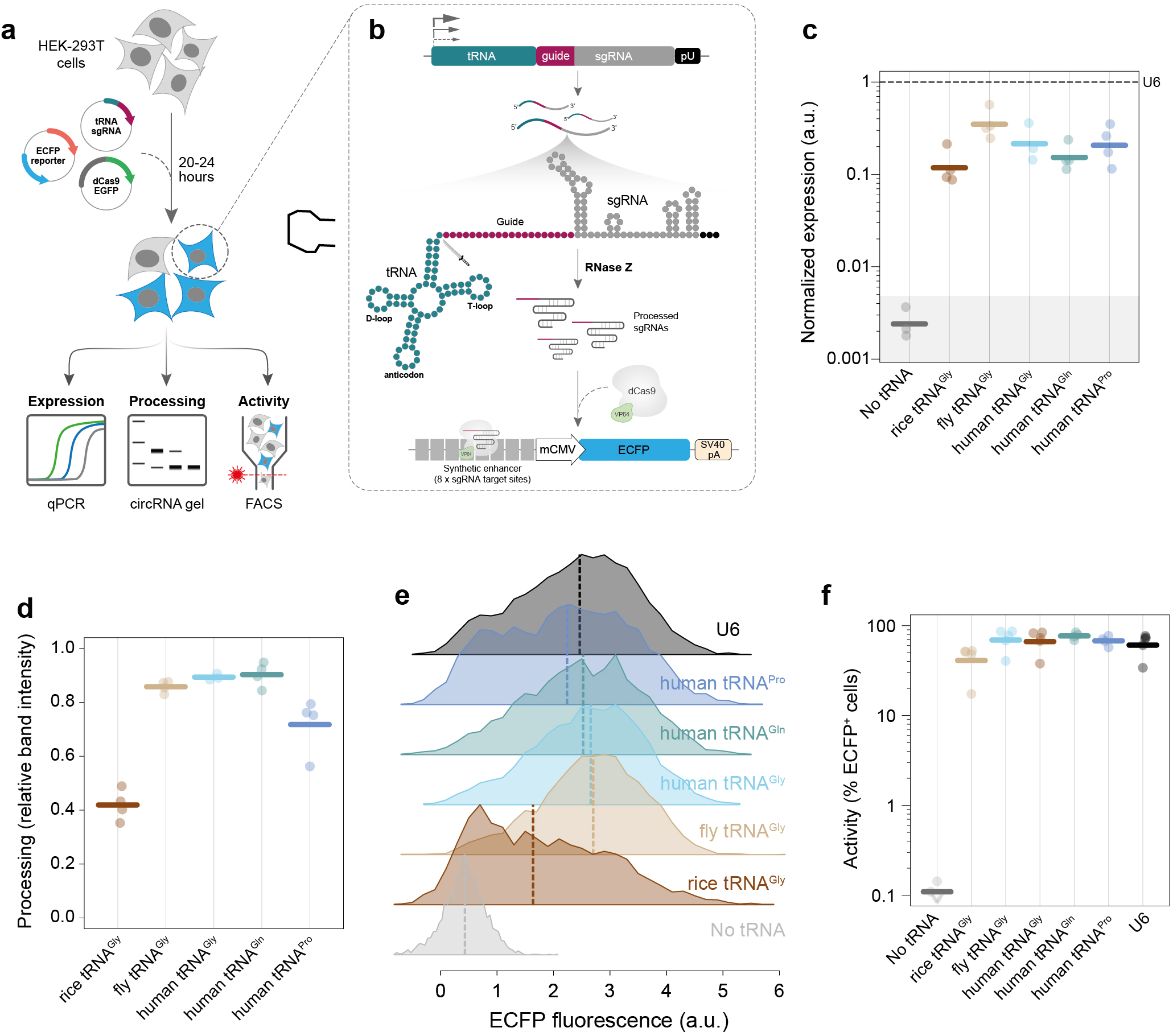
Wild type tRNAs display strong Pol-Ill promoter activity and gRNA production. **(a)** Experimental strategy for testing functional gRNA production downstream of tRNAs. **(b)** Schematic diagram of molecular events occurring in cells transfected with tRNA-sgRNA constructs. **(c)** gRNA expression as measured by qPCR relative to Cas9 and U6 (ΔΔCt) (n = 4 independent experiments, n = 3 for no promoter control; red line = gRNA levels for U6). Shaded area represents the 75% credible mass (BEST test) for the no promoter control. **(d)** 3’ processing ability of each wild-type tRNA tested. Efficiency represents the ratio of band intensity between the unprocessed and processed bands on a 2% agarose gel following RNA circularization and nested RT-PCR (thick lines = mean values). **(e)** Representative flow cytometry histograms of reporter levels downstream of U6 and various tRNA promoters. Values represent asinh(ECFP_fluoresence_/150) (thick lines = median fluorescent intensities), (f) Percent reporter ECFP^+^ cells within transfected cells (thick lines = geometric mean values; n=5 independent experiments; hollow downward triangles = points at or below the limit of detection).

**Supplementary Figure 2.**
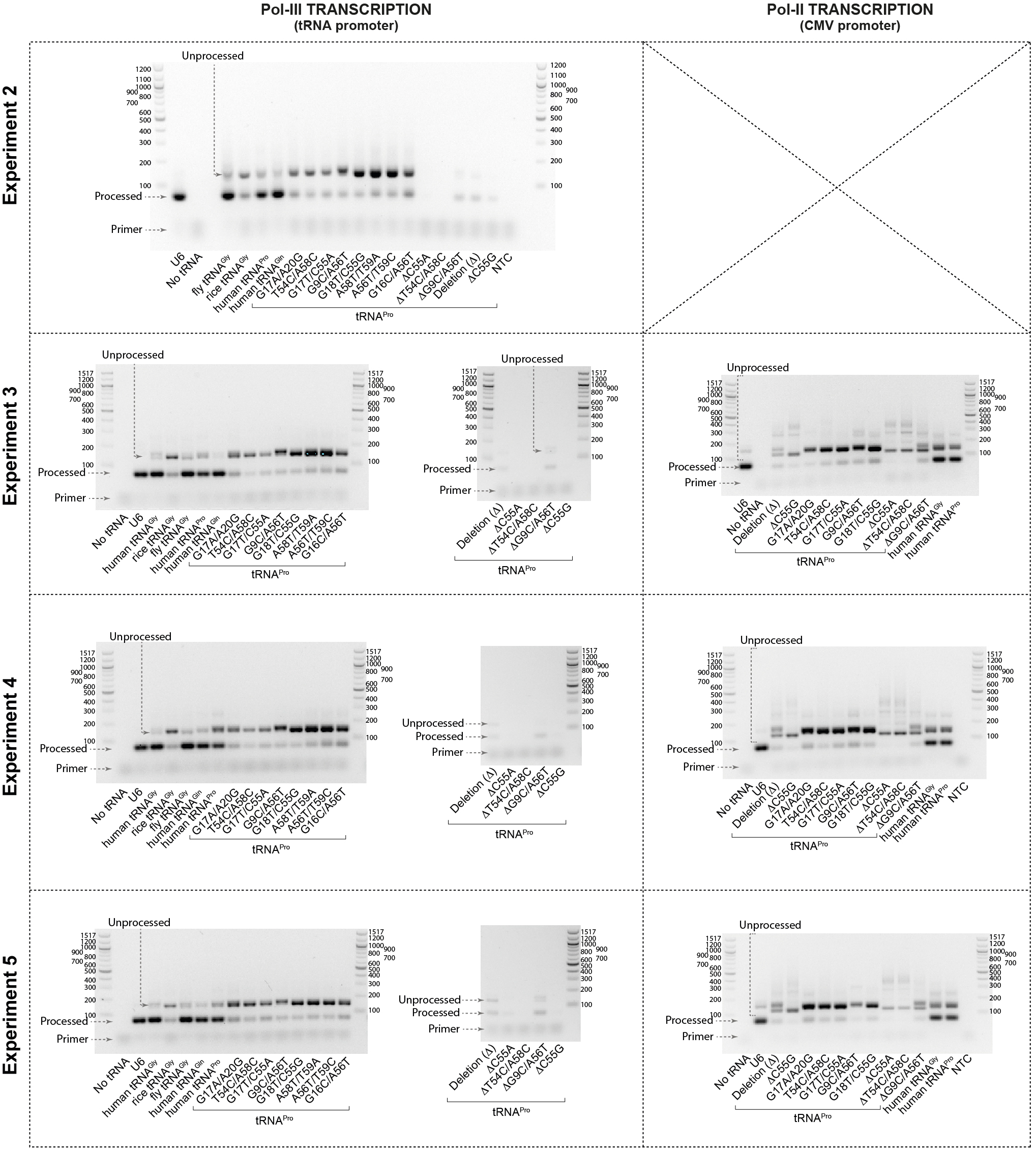
Agarose gel images used to determine processing efficiency. Gel images are shown for each experiment in which the RNA circularization assay was performed. Images from constructs with no external promoter are shown in the left column and those which had a Po1-II (CMV) promoter on the right. For Pol-II promoter containing assays RNA was first dephosphorylated prior to circularization. Size in base pairs (bp) are shown for all ladders. The approximate location of successfully processed bands, unprocessed bands, and residual primer are noted on each gel. Saturated pixels (where present) are shown in green.

**Supplementary Figure 3.**
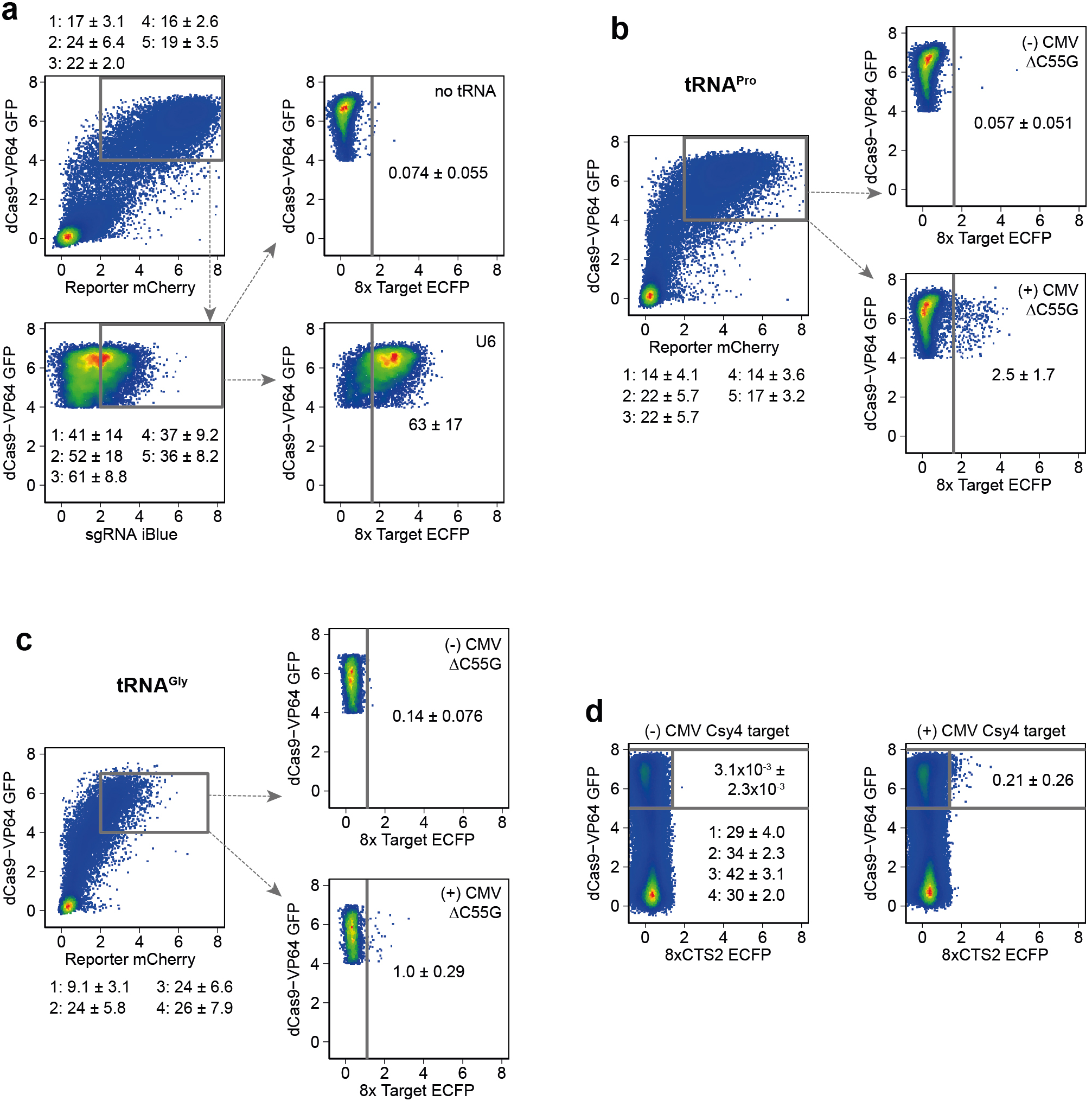
Representative gating strategies used in the manuscript. **(a)** Gating hierarchy used in all tRNA^Pro^ promoter testing experiments. Axes represent the asinh(ECFP_fluoresence_/150) of the fluorescence measurement (similar to a biexponential transformation). Values shown are mean ± s.d. for each experiment for the first two gates, and the mean ± s.d. across experiments for the final gate for the two examples shown. Plasmids expressed EGFP (dCas9-VP64), mCherry (in the reporter backbone), or iBlue (in the tRNA-sgRNA plasmids). Final ECFP gating was performed on cells positive for all requisite plasmids. In some cases, a maximum threshold was set on EGFP fluorescence where spillover would have been an issue. **(b, c)** Example of gating strategy for ON-OFF rates used for tRNA^Pro^ **(b)** and for tRNA^Gly^ **(c).** Mean ± s.d is shown per experiment for the first gate, and for the subsequent example final gates the mean ± s.d. across replicates is shown. In these instances, iBlue gating was not used in the gRNA plasmids as preliminary experiments suggested a crosstalk with the CMV promoter. **(d)** Gating strategy for alternative methods of gRNA processing. In this case only the EGFP on dCas9-VP64 was retained as other colors conflicted with some used in the alternative strategies. Mean ± s.d. % EGFP^+^ are shown for each experiment. Mean ± s.d. % ECFP^+^ (of EGFP^+^) are shown across replicates for the selected examples.

**Supplementary Figure 4.**
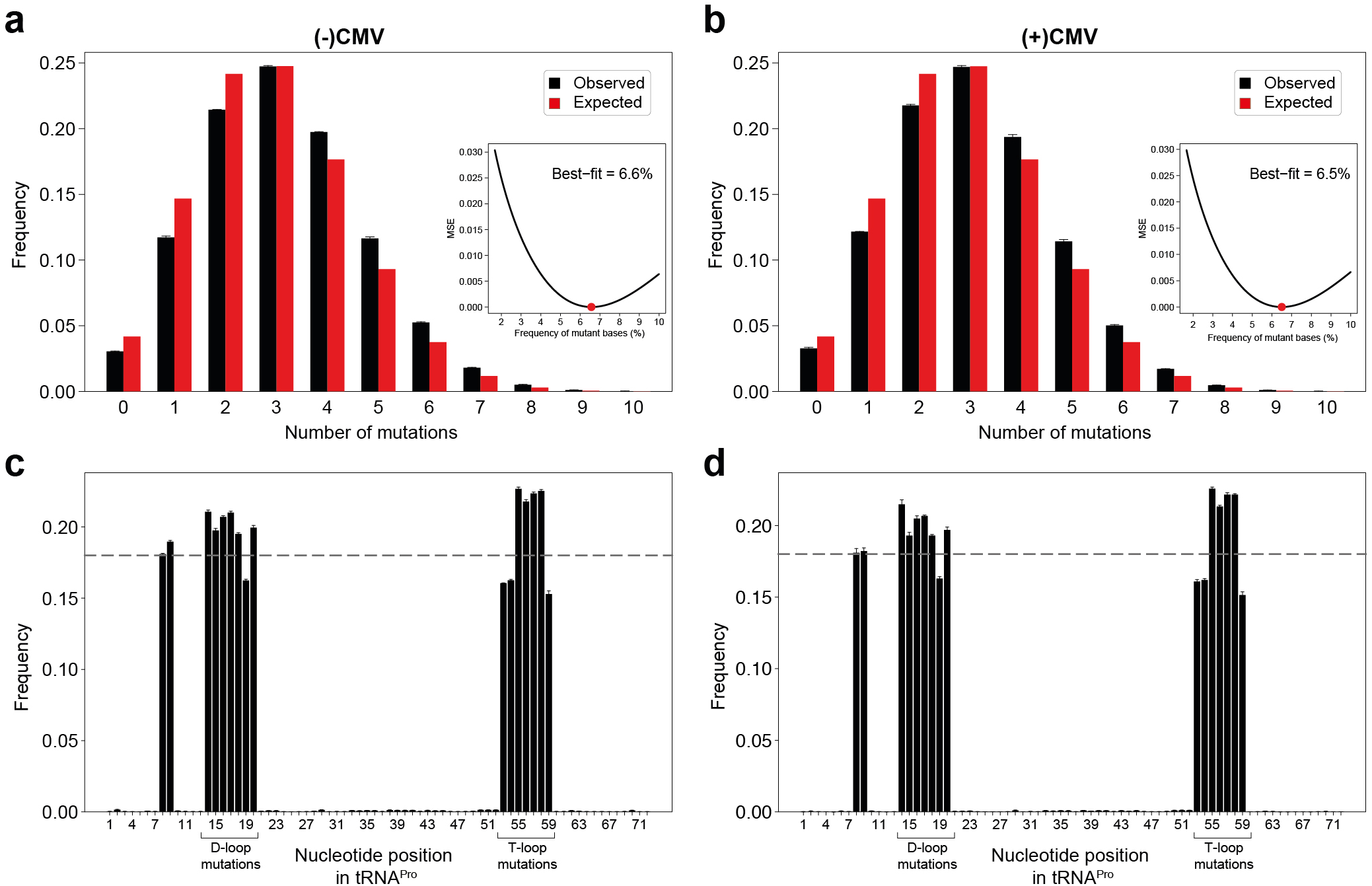
Mutation frequency and location in the tRNA screening libraries. **(a, b)** Frequencies for each number of mutations observed in the plasmid library without **(a)** and with **(b)** CMV promoters. The expected frequency based on the 6% of each non-WT base in the ordered oligo pool is shown in red. Black bars represent the frequency across three biological replicates for each library (mean ± s.d.). Insets show the mean-squared error (m.s.e.) of the measured mutational frequency compared to expected with differing frequencies of non-WT bases. Optimal fits are shown by red points and indicated in text. Best fits (minimum m.s.e.) are estimated at 6.6% and 6.5% without and with CMV promoters respectively, which closely match the 6% in the ordered pool. These results suggest that the diversity was not bottlenecked at any point in the protocol. **(c, d)** The frequency of mutations by location in the tRNA is shown for plasmid libraries without **(c)** or with **(d)** CMV promoters (n = 3 biological replicates, mean ± s.d.). These results confirm that mutant bases are specific for the targeted nucleotides. The expected overall mutation rate of 18% (given 6% of each non-WT base) is shown as a dotted line.

**Supplementary Figure 5.**
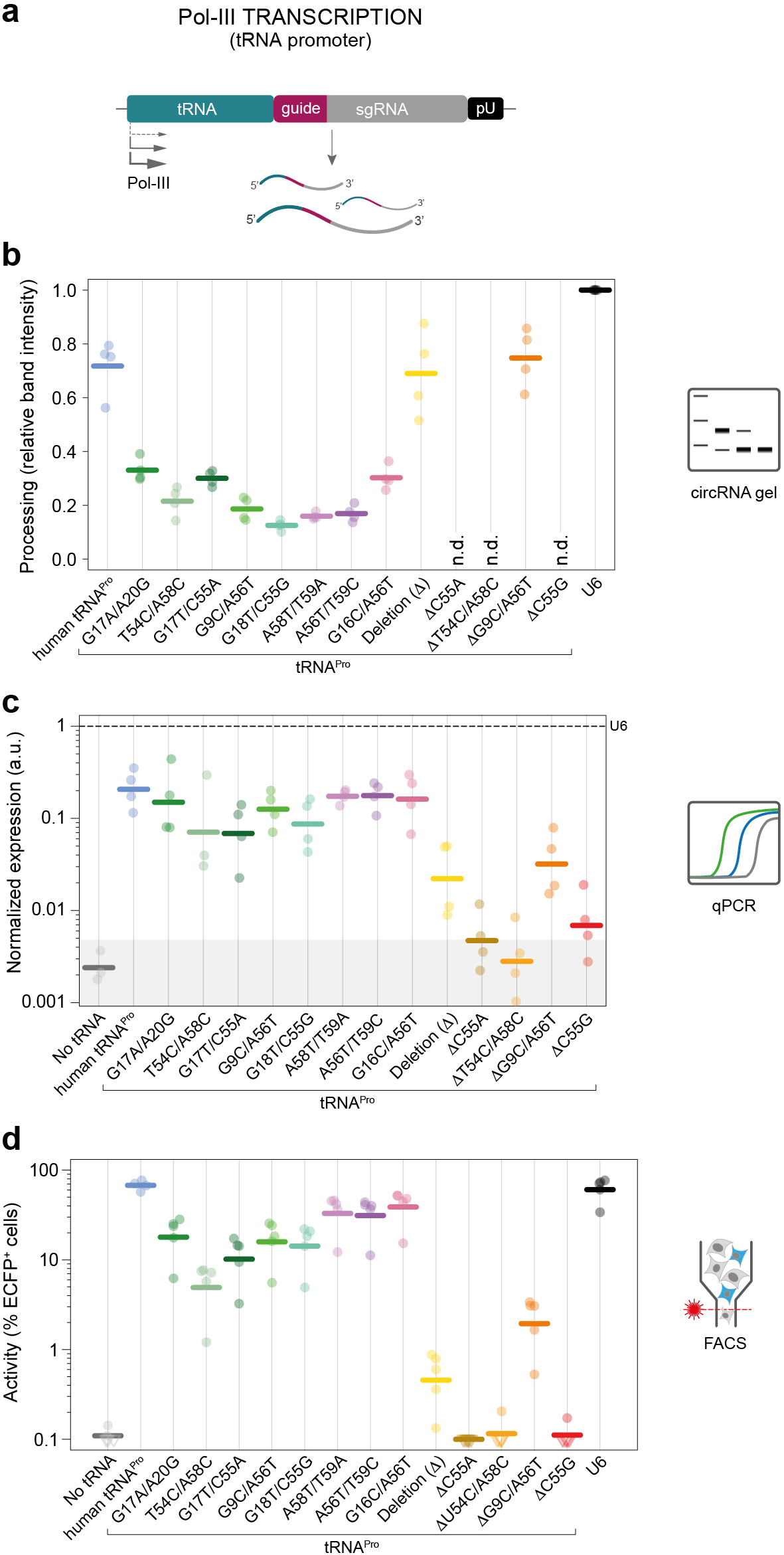
Characterization of promoter, 3’ processing and background gRNA production of engineered tRNA scaffolds. **(a)** Constructs and resulting RNA used for testing Pol-III promoter activity, **(b)** 3‘ processing ability of each mutant tRNA tested (n = 4 Independent experiments). Wild-type tRNA^Pro^ and U6 controls are shown for reference from Supplementary Fig. 1. Efficiency represents the ratio of band Intensity between the unprocessed and processed bands on a 2% agarose gel following RNA circularization and nested RT-PCR (thick lines = mean values; ND = variants with no detectable bands). All double-mutants showed some loss of 3’ processing compared to wild-type > 95% probability (paired BEST test). **(c)** gRNA expression as measured by qPCR relative to Cas9 and U6 (AACt) (n = 4 independent experiments, n = 3 for no promoter control; dashed line = gRNA levels for U6) (U6, ‘no tRNA, human tRNA^Pro^ and tRNA^Gly^ are shared with Supplementary Fig. 1c). Shaded area represents the 75% credible mass (BEST test) for the no promoter control. The T54C/A58C, G17T/C55A and G18T/C55G double-mutants predicted to decrease promoter activity (green tones) had a >80% probability, and all ΔtRNA scaffolds showed >95% probability of decreased promoter activity compared to wild-type (paired BEST tests), **(d)** Percent reporter ECFP^+^ cells within all transfected cells (thick lines = geometric mean values; each point = Independent experiment; n = 4-5). U6, wild-type tRNA^Pro^, and no promoter controls are shared with Fig. 2d, e and Supplementary Fig. 1 e, f. (hollow downward triangles = points at or below the limit of detection). Double-mutants affecting only processing (pink tones) had somewhat decreased functional activity compared to wild-type >99% probability, while those affecting promoter activity were even lower with >99% probability of decrease compared to all processing double-mutants (paired BEST tests).

**Supplementary Figure 6.**
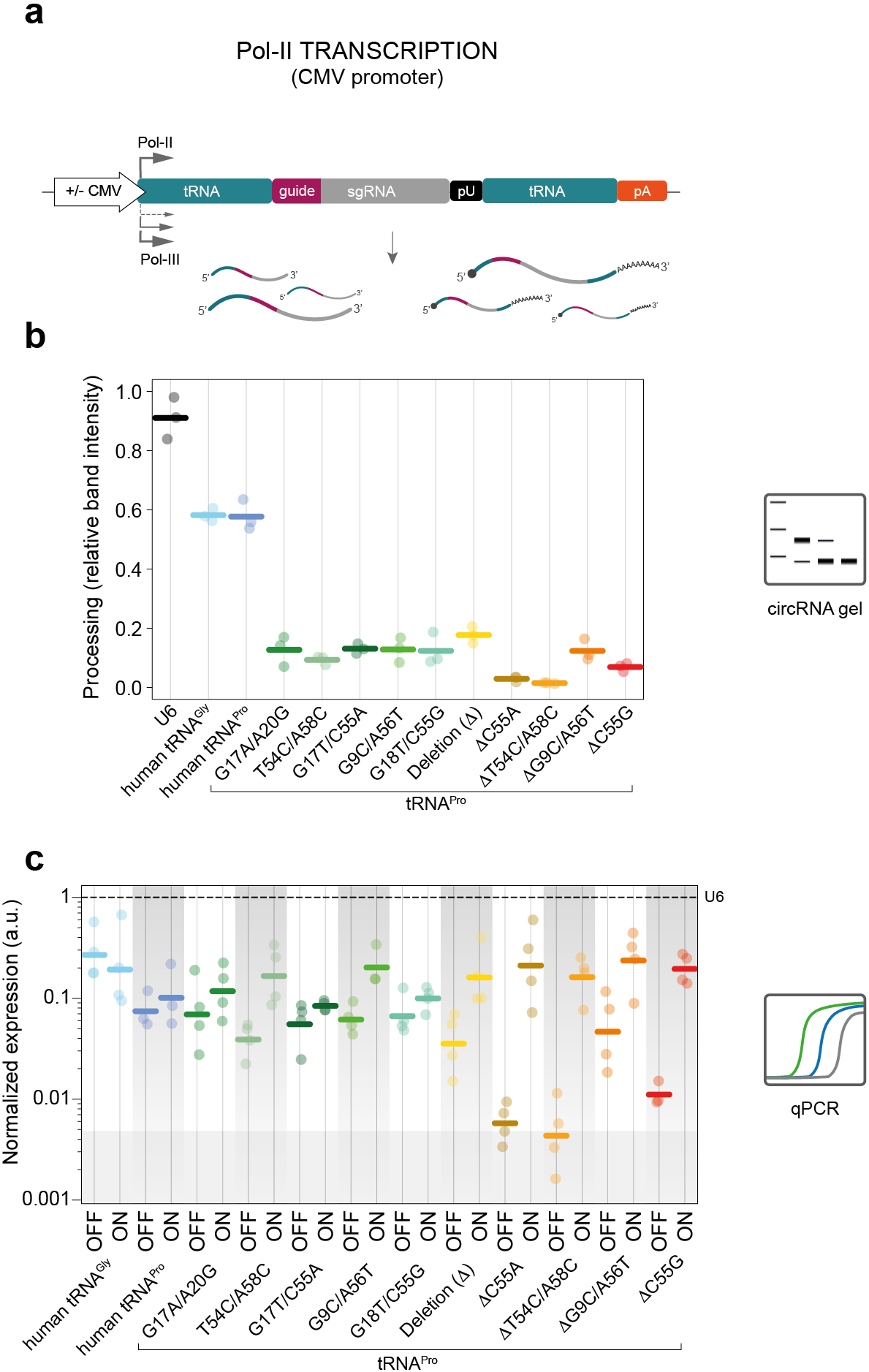
Characterization of engineered tRNA scaffolds overall processing and ON/OFF ratios in the presence and absence of Pol-II promoter. **(a)** Diagram of constructs used to test ON/OFF ratios in presence and absence of a Pol-II promoter (shared with Fig. 2c). **(b)** Overall processing ability of each mutant tRNA tested (n = 4 independent experiments). Efficiency represents the ratio of band intensity between the processed band and all other bands in the lane on a 2% agarose gel following decapping, RNA circularization and nested RT-PCR (thick lines = mean values), **(c)** gRNA expression for each mutation in the presence (ON) and absence (OFF) of a Pol-II promoter (CMV) as measured by qPCR relative to Cas9 and U6 (ΔΔCt) (n = 4 independent experiments, n = 3 for no promoter control; dashed line = gRNA levels for U6) (U6 and ‘no tRNA are shared with Supplementary Fig. 1c and 5c). Shaded area represents the 75% credible mass (BEST test) for the no promoter control.

**Supplementary Figure 7.**
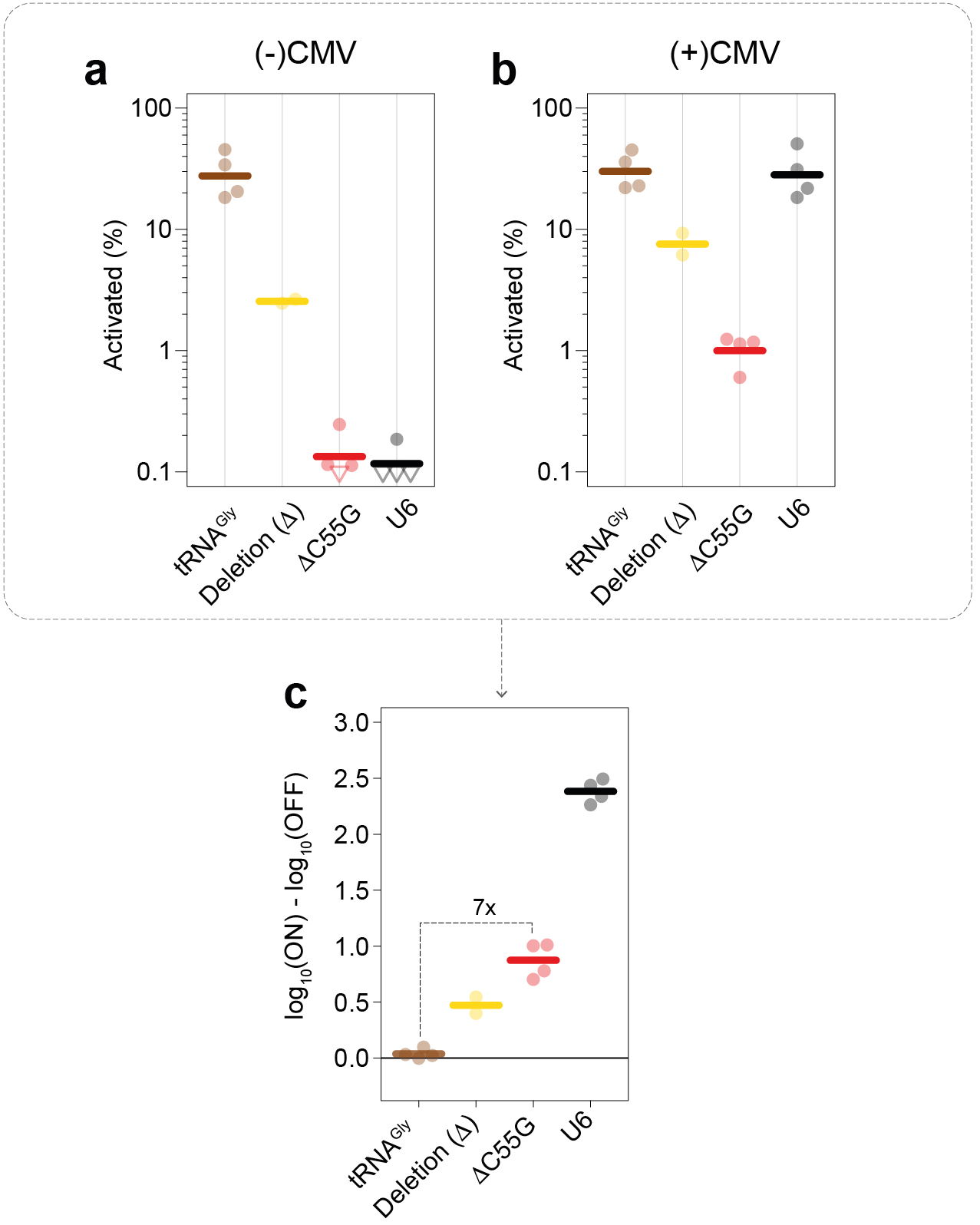
The new engineered scaffold is generalizable to other human tRNAs. **(a, b)** Percentage reporter ECFP^+^ cells within transfected cells for the tRNA^Gly^ constructs with **(a)** and without **(b)** CMV promoters. Points at or below the limit of detection are shown as hollow downward triangles, **(c)** Log_10_(% ECFP^+^ cells) in the ON condition (with CMV promoter) compared to OFF condition (no CMV promoter) for the tRNA^Gly^ constructs (n = 2-4 independent experiments). Thick lines represent mean values.

**Supplementary Figure 8.**
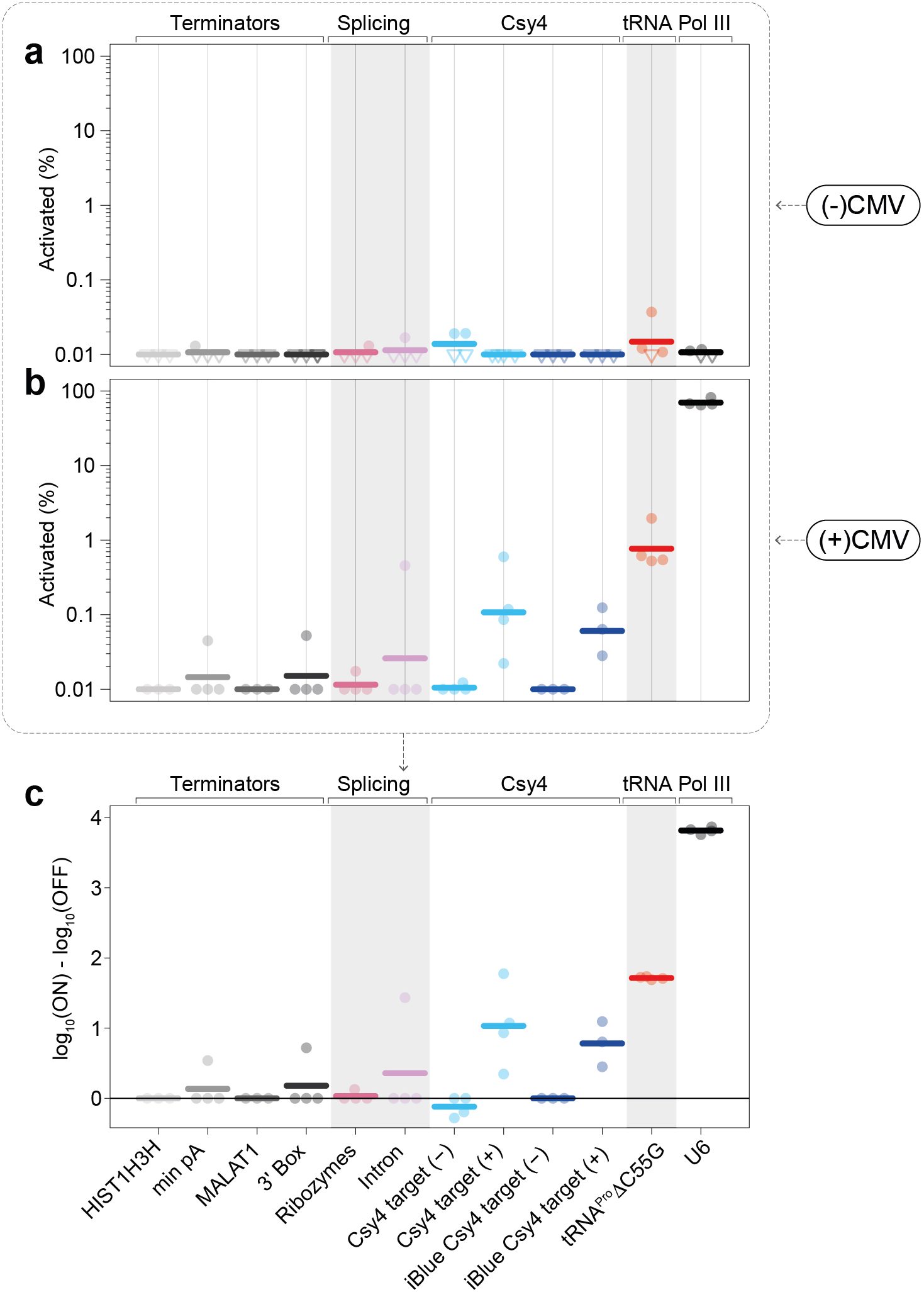
Comparison of top performing tRNA^Pro^ ΔC55G engineered variant with other reported systems for specific Pol-II gRNA expression. **(a, b)** Percent reporter ECFP^+^ cells within transfected cells for the OFF state (no CMV promoter) **(a)** and the ON-state (with CMV promoter) **(b).** Points at or below the limit of detection are shown as hollow downward triangles. **(c)** Log_10_(% ECFP^+^ cells) in the ON condition (with CMV promoter) compared to OFF condition (no CMV promoter). Thick lines represent mean values (n = 3-4 independent experiments). Terminators refer to the use of alternative termination sequences to the standard poly-A and include the histone 1h3h terminator, a minimal poly-A sequence^5^, the MALAT 1 terminator^5^, as well as the U1 snoRNA3’ box^5^. The ribozyme scaffold contains a gRNA flanked by hammerhead and HDV ribozymes^7^. The intron system contains a gRNA embedded within an artificial intron inside the mKate fluorphore^6^. Cys4 refers to a construct in which the gRNA is flanked by minimal Csy4 targets^19^ either on its own in a transcript, or in the 3’ UTR of an iBlue transcript, in the presence (+) or absence (-) of Csy4 protein. Csy4 constructs had slightly lower functional gRNA compared to AC55G with 88.7% and 90.7% probability of decrease in paired BEST tests.

## ONLINE METHODS

### Rationale for tRNA variant screening

Quantitative PCR (qPCR) of gRNAs placed downstream of native tRNAs revealed robust constitutive guide production in the absence of external promoters in all cases, albeit at lower overall levels than U6-driven gRNA expression (≥92% probability, Bayesian Estimation Supersedes the t-test “BEST” test^20,21^, Supplementary Fig. 1c). This is consistent with findings that tRNA promoters appear to be slightly less efficient than U6 for gRNA production^12^. All human and fly tRNAs tested showed very efficient 3’ processing activity as measured using a modified circularization assay^9^ (Supplementary Fig. 1d, 2). Importantly however, when tested in a gRNA functional reporter assay^4,22^, all tRNAs with the exception of rice tRNA^Gly^ enabled efficient transcriptional activation at levels equivalent to U6, in the absence of any additional Pol-II or Pol-III promoters (Supplementary Fig. 1e, f, 3a).

Remarkably, fly and rice tRNA^Gly^ showed promoter activity, processing ability, and functional gRNA production in human cells, albeit the values displayed by rice tRNA^Gly^ were reduced compared to human tRNA^Gly^ (BEST probabilities of decreased effect 89%, Supplementary Fig. 1 and 3a). These results reflect the strong conservation of tRNA systems across kingdoms and suggest that the principles described in this study will likely be applicable to a number of model organisms.

### Cloning and construct assembly - standard ligations and transformations

All restriction enzyme digestions were performed in suggested buffers either as double digestions (if compatible) or sequential digestions as appropriate. In the case of sequential digestions, a Qiaquick PCR purification (Qiagen) with 30 μl elution volume (Buffer EB) was used to re-purify between digestions. Following digestion, vectors were treated with 5 U of Antarctic Phosphatase for 30 minutes at 37°C (NEB). All vectors were then gel purified using a Qiagen gel extraction kit as per manufacturer instructions but with Qiagen MinElute PCR purification columns substituted for Qiaquick columns and a final elution volume of 15 μl buffer EB. Inserts were purified prior to ligation with either a standard Qiagen gel extraction protocol, or a Qiaquick PCR purification (Qiagen) as appropriate. All standard ligations were performed using 200 or 400 U T4 DNA ligase in T4 DNA ligase buffer (NEB) with approximate insert to vector ratios between 1:1 and 1:10. Ligation reactions were incubated at room temperature for 5-30 minutes (5-10 minutes for single insert ligations, 20-30 minutes for ligations with more inserts). Ligase was then heat inactivated by 10 minute incubation at 80°C. Ligation reactions were then cooled on ice for 2 minutes, then 1-3 μl added to 10-50 μl Subcloning Efficiency™ DH5α™ Competent Cells (Thermo Fisher Scientific) with a maximum ligation to bacteria ratio of 1:10. Cells were then incubated on ice for 15-30 minutes, heat shocked at 42°C for 20 seconds in a water bath, then 200-500 μl S.O.C. medium (homemade) added. Cells were then incubated for 40 minutes to 1 hour at 37°C with shaking and 200 μl plated onto LB Agar plates with 100 μg/ml Ampicillin. Plates were incubated at 37 °C overnight. Individual colonies were then picked into 5-7 ml LB + 100 μg/ml Ampicillin (Sigma) and incubated again overnight at 37°C with shaking. Plasmids were then purified using Qiaprep Spin Miniprep columns (Qiagen). Finally, plasmids were verified by appropriate diagnostic digests and sequencing with appropriate primers (generally one of pBR322_ori-F: CACCTCTGACTTGAGCGTCG, AmpR-R: GGTTATTGTCTCATGAGCGG, SV40polyA-R: TACTCGAGGGATCCTTATCGATTTTACC, or forUAS-F: CCAATCTCGAGGAGGCTAGGGATGAAGAATAAAAG) using Eurofins Genomics sequencing service.

### Cloning and construct assembly - PCR amplifications for cloning

For all PCR amplifications used in cloning, Phusion^®^ High-Fidelity PCR Master Mix with GC Buffer (New England BioLabs, NEB) was used for amplification with 500 nM of each primer (primers are as indicated for individual reactions). Amplification conditions were as follows with ‘*’ indicates optimal annealing conditions as determined by NEB Tm calculator.

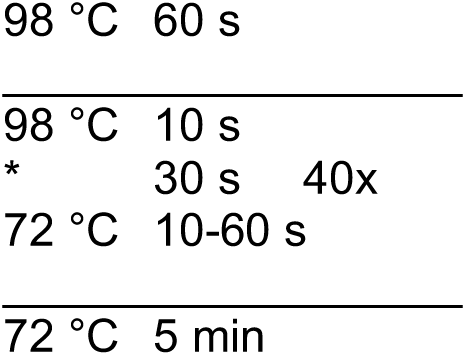

### Cloning and construct assembly - oligonucleotide phosphorylation and annealing

For all oligo annealing reactions 10 μl containing 10 μM each of the forward and reverse oligos, 1x T4 DNA ligase buffer, and 5 units of T4 Polynucleotide Kinase (NEB) were incubated at 37°C for 30 minutes, ramped to 95°C for 5 minutes then cooled to 25°C at a rate of 0.1°C/s. The annealed oligos were then used as inserts for cloning reactions where applicable.

### Cloning and construct assembly - backbone creation

An initial vector was created by three insert restriction cloning using a backbone containing a pBR322 origin with ampicillin resistance and adjacent SphI and SbfI restriction sites. To generate the first insert, we first amplified the poly-A signal and pause site out of the 8xCTS2-mCMVp-EYFP construct (REF^22^) (Forward: GCTAGCGGTACCGGTACTTGGAGCGGCCGC, Reverse: GGCGCCGGTACCCGATAGAGAAATGTTCTGGCACCTG). The amplified fragment was purified by Qiaquick PCR purification (Qiagen) and incubated with 5 units of standard Taq polymerase in standard Taq buffer with 2.5 mM dATP (NEB) for 15 minutes at 72°C to add A overhangs. This was then TA cloned using a TOPO® TA Cloning® Kit (Thermo Fisher Scientific). Resulting clones were then reamplified (Forward: TTTTGCTCACATGTGCATGCGGAGCGGCCGCAATAAAATATCT, Reverse: CTAGGGCGCTGGCAAGTGTA) and the resulting product digested with SphI and KpnI to obtain the first insert. A modular scaffold containing multiple cloning sites and a minimal CMV promoter (mCMV) promoter was ordered from IDT as a gBlock (CAGTGCAAGTGCAGGTGCCAGAACATTTCTCTGGGCCCGCCCGGTACCTTCTAGAGGCTAGGGATGAAGAATAAAAGGGGCGCGCCCCTAGGGAGCGGGGGGCTATAAAAGGGGGTGGGGGCGTTCGTCCTGCTATCTAGCGTCGCGTTGACCGAGCTCATCGGTGCATGGGTGGTTCAGTGGTAGAATTCTCGCCTGCCACGCGGGAGGCCCGGGTTCGATTCCCGGCCCATGCAGGAGACGGACGTCTCCGTTTTAGAGCTAGGCCAACATGAGGATCACCCATGTCTGCAGGGCCTAGCAAGTTAAAATAAGGCTAGTCCGTTATCAACTTGGCCAACATGAGGATCACCCATGTCTGCAGGGCCAAGTGGCACCGAGTCGGTGCAACAAAGCACCAGTGGTCTAGTGGTGGAATAGTACCCTGCCACGGTACAGACCCGGGTTCGATTCCCGGCTGGTGCAGGCCGGCCGC). This was amplified (Forward: CAGTGCAAGTGCAGGTGCCA, Reverse: GCGGCCGGCCTGCAC), A overhangs added as for insert one, and TA cloned using a TOPO^®^ TA Cloning^®^ Kit (Thermo Fisher Scientific). Resultant TOPO clones were digested with KpnI and FseI to obtain the middle insert. The final insert, the SV40 polyA sequence, was amplified from the 8xCTS2-mCMVp-ECFP construct (REF^22^) (Forward: CCCGGCTGGTGCAGGCCGGCCGCTTC, Reverse: GCCACCTGACGTCCCTGCAGGCTCGAGGGATCCTTATCGATTTTACC) followed by digestion with FseI and SbfI-HiFi (both from NEB). This resulted in the minimal backbone used for downstream cloning, *Backbone 1*. Finally, to allow gating for cells which received the gRNA plasmids, the SV40 promoter-iBlue-SV40 polyA site was amplified out of the U6-CTS2 construct (REF^22^) (Forward: ACGGAGACGTCGAATGTGTGTCAGTTAGGGTG, Reverse: TACGTCTCGAGTTAGCTCACTCATTAGGCAC), digested with XhoI and AatII and inserted into the same sites in *Backbone 1* to generate *Backbone 2*.

### Cloning and construct assembly - 8xCTS2 reporter modification

To allow the functional analyses to be performed only on cells which had received all necessary plasmids, we inserted a SV40 promoter-mCherry-SV40 polyA cassette into the backbone of the 8xCTS2-mCMVp-ECFP construct (REF^22^) between the PciI and SalI sites. The insert was amplified from an available plasmid using primers which added the PciI and SalI sites respectively (Forward: TGCTTACATGTGGAATGTGTGTCAGTTAG, Reverse: CTAACCTCGAGCAAGCTCTAGCTAGAGGTCG).

### Cloning and construct assembly - tRNA promoter testing

To create the constructs used in testing the Pol-III promoter activity of tRNA, 3 - 4 insert standard cloning was performed. The tRNAs comprised 1 or 2 pairs of (separately) annealed oligos as listed in Supplemental Table 1. In cases with a mutation in one part but not the other, the wild-type oligo pair was used for the opposite side insert. The CTS2 guide sequence was another pair of annealed oligos (Supplemental Table 1). All annealed oligo pairs had unique 4 bp overhangs on each side to allow scar-free ligation to their partners. Finally, the single guider RNA (sgRNA) was amplified from pX330-U6-Chimeric_BB-CBh-hSpCas9 (Addgene plasmid # 42230, a gift from Feng Zhang) using forward primer, CGAGAAGACCTGTTTTAGAGCTAG, and reverse primer GAAGCGGCCGGCCAAAAAAGCACCGACTCG, followed by a sequential digest with BpiI (Thermo Fisher Scientific) and FseI (NEB). These inserts were placed into *Backbone 2* between the SacI and FseI sites (NEB). In some cases, the minimal CMV was then replaced using AvrII and SacI-HF (both from NEB) with the annealed oligo set ‘noPromoter’. This was not done for the D-loop anti-codon deleted tRNA^Pro^ and the ΔC55G tRNA^Pro^, as in functional tests it was not shown to make a difference.

### Cloning and construct assembly - Pol-II construct cloning

Cloning for constructs to test the effects of Pol-II promoters was done in a 2 ((-)CMV) or 3 ((+)CMV) stages. First, the existing tRNA variant was amplified using appropriate primers (Supplemental Table 2) with an added FseI restriction site on each side. This product was then cloned into the FseI site of the parental plasmid (containing a tRNA-CTS2 sgRNA) located between the sgRNA and SV40 polyA site. Following screening for correct orientation (by sequencing), the SV40 promoter-iBlue-SV40 polyA cassette was removed by AatII+XhoI digestion (NEB) and replaced with the annealed oligo set ‘iBlueRemover’. This was done in all constructs aside from the D-loop/Anticodon deleted tRNA^Gly^ and the ΔC55G tRNA^Gly^ as in initial tests Pol-II specific activation was anticorrelated with iBlue levels, suggesting possible crosstalk with the Pol-II tRNA cassette, although the functional activation levels achieved ignoring the iBlue did not appear to be affected. This step yielded the (-)CMV constructs. Finally, a CMV promoter amplified from a microRNA reporter plasmid (Michaels et al. in preparation, Forward: ACGTTGGCGCGCCCGAGCATTAGTTCATAGCCCATATATGG, Reverse: GCAACGAGCTCGACCGGTGGATCTG) was inserted into the AscI-SacI site in the (-)CMV construct (replacing the minimal CMV or ‘noPromoter’ region).

### Cloning and construct assembly - alternative processing strategies

For the intronic guide constructs, we ordered a gBlock (below) containing mKate with an intronic sgRNA backbone and a cloning site to insert the gRNA sequence based on REF^6^. This was amplified to add in restriction sites for cloning into *Backbone 1* (Forward: AAGTAGAGCTCGCCACCATGGTGTCTAAGG, Reverse: AATGAGGCCGGCCTCAATTAAGTTTGTGCCCCA). This was then cloned between the SacI and FseI sites of *Backbone 1* to give an intermediate construct for the (-)CMV version. Simultaneously, the same CMV promoter amplicon from the Pol-II constructs was co-inserted into *Backbone 1* together with the mKate intronic gRNA construct between the AscI and FseI sites of *Backbone 1* to give the (+)CMV intermediate. Annealed oligos (‘intronic CTS2’) were then inserted into the gRNA cloning site of each of these to clone the final intronic gRNA constructs.

To clone the ribozyme release system, a three-insert cloning between the SacI and FseI sites of the (+)CMV and (-)CMV intronic gRNA constructs was performed (removing the complete mKate/intronic guide). The first insert was a set of annealed oligos, ‘HH_CTS2’, containing the hammerhead ribozyme from REF^7^ specific for CTS2 (the first 6 bases of the ribozyme complementary to the first 6 bases of the guide) with the CTS2 guide. This pair contained 4 bp overhangs compatible with SacI and the sgRNA. The second insert was the sgRNA 2.0 with a BsmBI cloning placeholder. This was amplified from *Backbone 1* (Forward: TGCCAGAAGACATGGAGACGGACGTCTCCGTTTTAG, Reverse: GAGGCGAAGACTAGCACCGACTCGGTGCCA) and digested with BsmBI and BpiI to give compatible sticky ends. The final insert was the HDV ribozyme from REF^7^. This ribozyme sequence was ordered as an oligo from IDT (GGCCGGCATGGTCCCAGCCTCCTCGCTGGCGCCGGCTGGGCAACATGCTTCGGCATGGCGAATGGGAC) and amplified to add a BpiI site to the front such that the digested product had sticky ends compatible to the sgRNA and FseI site to the back (Forward: GAGGCGAAGACTAGTGCGGCCGGCATGGTCCC, Reverse: AAGTAGGCCGGCCGTCCCATTCGCCATGCC). This was sequentially digested with BpiI and FseI to give the final fragment.

For all alternative transcriptional terminators, the (+)CMV intronic gRNA construct was used as a vector. Inserts were then placed between the AgeI and XhoI site for the (+)CMV terminator constructs (leaving the initiator consensus of the CMV promoter immediately upstream of the guide while removing the SV40 polyA), and between the AscI and XhoI sites for the (-)CMV terminator constructs (removing the CMV and SV40 poly A entirely). In all cases this was done by 2-insert cloning. The first insert contained the gRNA amplified from the ribozyme construct using primers which added BpiI sites on either end. For the (-)CMV constructs the BpiI site in the forward primer (ATTTAGAAGACAACGCGCCGTAAGTCGGAGTACTGTCCTGTTTTAGAGC) created a sticky end compatible with AscI, and for the (+)CMV constructs the BpiI site in the forward primer (ATTTAGAAGACAACCGGTGTAAGTCGGAGTACTGTCCTGTTTTAGAGC) created a sticky end compatible with AgeI. In both cases these shared a common reverse primer (GAGGCGAAGACTAGCACCGACTCGGTGCCA). The second insert contained the terminators. This was made up of annealed oligos with 4 bp overhangs compatible with the gRNA at the 5’ end and with the vector XhoI site on the 3’ end for the minimal polyA and HIST1H3H terminators. For the MALAT1 and 3’ Box terminators the sequences were ordered from IDT (below) then amplified with BpiI flanked primers to yield compatible sticky ends (MALAT1 Forward: TGCCAGAAGACATGTGCGATTCGTCAGTAGGGTTGTAAAGG, MALAT1 Reverse: TGCCAGAAGACATTCGAGAAGCAAAGACACCGCAGG; 3’ Box Forward: ATTTAGAAGACAAGTGCGGATCCACTTTCTGGAGTTTCAAAAG, 3’ Box Reverse: ATTTAGAAGACAATCGAGTTAAGACGCCAACCAAG).

The basic Csy4 constructs were cloned as a single insert into the SacI FseI site of the intronic gRNA constructs. The insert containing the gRNA was amplified from the HIST1H3 terminator construct using primers which added the minimal Csy4 sequence (CTGCCGTATAGGCAGC) and SacI and FseI sites respectively (Forward: TGCCAGAGCTCCTGCCGTATAGGCAGCGTAAGTCGGAGTACTGTCCTGTTTTAGAGC, Reverse: TGCCAGGCCGGCCGCTGCCTATACGGCAGGCACCGACTCGGTGCCA). The iBlue-Csy4-gRNA constructs were made in a two-insert cloning between the AscI and FseI sites of the intronic guide constructs. The first insert was amplified from a microRNA reporter plasmid (Michaels et al. in preparation) and contained either the iBlue sequence alone (Forward: TGGATGGCGCGCCTAGTGAACCGTCAGATCCAC, Reverse: ACAGAGAGCTCAGTTATTAGGTCCCTCGACG), or a CMV promoter followed by iBlue (Forward: ACGTTGGCGCGCCCGAGCATTAGTTCATAGCCCATATATGG, Reverse: ACAGAGAGCTCAGTTATTAGGTCCCTCGACG). In both cases these were flanked by AscI sites and SacI sites to make the inserts compatible with the vector and same gRNA amplicon used for the basic Csy4 constructs (the second insert).

#### mKate intronic sgRNA2.0 gBlock

GCCACCATGGTGTCTAAGGGCGAAGAGCTGATTAAGGAGAACATGCACATGAAGCTGTACATGGAGGGCACCGTGAACAACCACCACTTCAAGTGCACATCCGAGGGCGAAGGCAAGCCCTACGAGGGCACCCAGACCATGAGAATCAAGGTGGTCGAGGGCGGCCCTCTCCCCTTCGCCTTCGACATCCTGGCTACCAGCTTCATGTACGGCAGCAAAACCTTCATCAACCACACCCAGGGCATCCCCGACTTCTTTAAGCAGTCCTTCCCTGAGGTAAGTGGTCCGGAGACGGACGTCTCCGTTTTAGAGCTAGGCCAACATGAGGATCACCCATGTCTGCAGGGCCTAGCAAGTTAAAATAAGGCTAGTCCGTTATCAACTTGGCCAACATGAGGATCACCCATGTCTGCAGGGCCAAGTGGCACCGAGTCGGTGCTACTAACTCGAGTCTTCTTTTTTTTTTTCACAGGGCTTCACATGGGAGAGAGTCACCACATACGAAGACGGGGGCGTGCTGACCGCTACCCAGGACACCAGCCTCCAGGACGGCTGCCTCATCTACAACGTCAAGATCAGAGGGGTGAACTTCCCATCCAACGGCCCTGTGATGCAGAAGAAAACACTCGGCTGGGAGGCCTCCACCGAGATGCTGTACCCCGCTGACGGCGGCCTGGAAGGCAGAAGCGACATGGCCCTGAAGCTCGTGGGCGGGGGCCACCTGATCTGCAACTTGAAGACCACATACAGATCCAAGAAACCCGCTAAGAACCTCAAGATGCCCGGCGTCTACTATGTGGACAGAAGACTGGAAAGAATCAAGGAGGCCGACAAAGAGACCTACGTCGAGCAGCACGAGGTGGCTGTGGCCAGATACTGCGACCTCCCTAGCAAACTGGGGCACAAACTTAATTGA sgRNA2.0, mKate, Splice Donor, Splice Acceptor, Polypyrimidine tract, Branch Point, Guide cloning site

#### MALAT1 terminator

GATTCGTCAGTAGGGTTGTAAAGGTTTTTCTTTTCCTGAGAAAACAACCTTTTGTTTTCTCAGGTTTTGCTTTTTGGCCTTTCCCTAGCTTTAAAAAAAAAAAAGCAAAAGACGCTGGTGGCTGGCACTCCTGGTTTCCAGGACGGGGTTCAAGTCCCTGCGGTGTCTTTGCTT

#### 3’ Box terminator

ACTTTCTGGAGTTTCAAAAGTAGACTGTACGCTAAGGGTCATATCTTTTTTTGTTTGGTTTGTGTCTTGGTTGGCGTCTTAA

### Cloning and construct assembly - tRNA screening constructs

Cloning for the single nucleotide variant tRNA libraries was performed in 3 (-)CMV or 4 (+)CMV stages. For the (+)CMV constructs, first the CMV promoter amplicon used for the Pol-II constructs was inserted between the AscI and SacI sites of one of the Pol-III tRNA constructs (tRNA-pol-III terminator, SV40 promoter-iBlue-SV40 polyA). The next stage (shared between the (+)CMV and (-)CMV libraries) involved a three-insert cloning. The first insert comprised a CTS2-sgRNA amplified from the Pol-III tRNA constructs with primers adding a SacI site to the 5’ end and a BpiI site to the 3’ end such that the overhang was complementary to the middle insert (Forward: AGCACGAGCTCGTAAGTCGGAGTACTGTCCTGTTTTAGAG, Reverse: TCGAAGACCCGCACCGACTCGGTGCC). The second insert was a pair of annealed oligos containing a pair of outward facing BpiI sites to allow downstream GoldenGate assembly (‘Bpil_placeholder’). This annealed oligo had 4 bp overhangs complementary to the overhangs created by the BpiI digestions of the inserts on either side. The final insert was also a CTS2-sgRNA amplified from the Pol-III tRNA constructs but with primers now adding a BpiI site to the 5’ end, retaining the pol-III terminator, and adding a FseI site to the 3’ end (Forward: TAAGAAGACTAGTAAGTCGGAGTACTGTCCTGTTTTAGAG, Reverse: GAAGCGGCCGGCCAAAAAAGC). Sequencing validation of the (-)CMV construct at this stage revealed a recombination event between the SV40 polyA downstream of the main construct and downstream of the iBlue (which is in the reverse orientation). This retained bidirectional polyA sites thus still allowing proper iBlue transcript formation, and the event does not affect the Pol-III transcribed region. Therefore, this construct was still retained for downstream cloning.

We next added in the flanking barcode sequences and buffers with a paired BsmBI site between them by BpiI-based GoldenGate assembly. To do so, we ordered a 116 bp barcode library ultramer from IDT (‘Barcode library’ sequence below). This was amplified in three independent PCR reactions with 100 pmol template per reaction (Forward: TAACGAGGCGAAGACTAGTGC, Reverse: GCAATGCCAGAAGACATTTAC). Following amplification products were gel purified using Qiagen gel extraction kit as per manufacturer instructions but with Qiagen MinElute PCR purification columns. These were then combined at equimolar ratios to give the final insert. This was done to minimize potential PCR error and sampling biases^23,24^. A total of 20 GoldenGate assembly reactions were then assembled and cycled as follows:

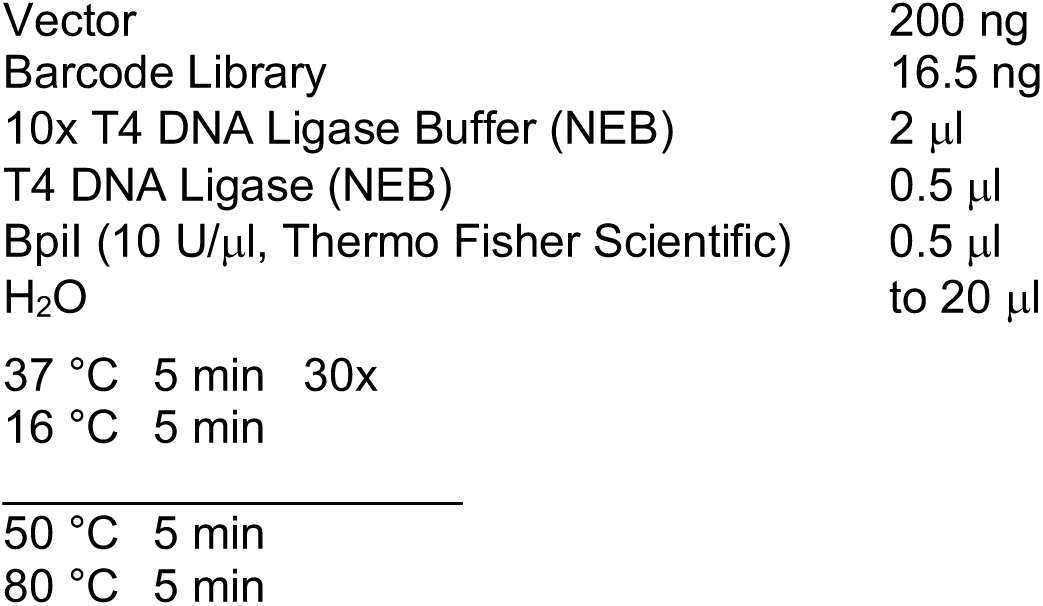

Following cycling all 20 GoldenGate reactions were pooled and co-purified by Qiaquick PCR purification (Qiagen) with a 30 μl elution volume. A total of three transformations were performed. For these, 5 μl of the purified GoldenGate reaction was then added to 30 μl of NEB 10-beta Competent E. coli (NEB) and transferred to an electroporation cuvette with a 1 mm gap (VWR). This was then electroporated on an Eppendorf Eporator^®^ with 1.6 kV over 5 ms. Volume was then immediately topped up to 1 ml with 37°C SOC (home-made) and transferred to a 1.5 ml microtube and incubated at 37°C with shaking for 1 hour. All 3 transformations were then combined and the total volume brought to 8 ml with warm SOC. This was used to inoculate two 245 mm dishes (Corning). 0.1 μl was also plated onto a 10 cm dish to allow an estimation of colony number. These were incubated for 16 hours at 32°C. Following incubation colonies were harvested by scraping using bacteria spreaders with LB washes. Harvested colonies were collected into a 50 ml Falcon tube (Thermo Fisher Scientific), spun down at 3000g, and the supernatant removed. Pellets were then weighed and split across a multiple Qiagen plasmid Midiprep columns and plasmids purified as per manufacturer protocols.

For the final stage of library cloning 36 μg of each vector from the previous stage was digested in a 360 μl reaction with a total of 120 U of BsmBI in NEBuffer 3.1 (both from NEB) for 2 hours at 55°C.

Following digestion, 60 units of Antarctic Phosphatase and associated buffer was added to the reaction and incubated for an additional 30 minutes at 37°C. Digestions were then gel purified with a Qiagen gel extraction kit as per manufacturer instructions. Inserts were prepared by triplicate PCR amplification (Forward: TAACGTCTCTCGTCGGCTCG, Reverse: TAACGTCTCAGGGCTCGTCC) from the tRNA variant library oligo (below), gel purification, and equimolar pooling as was done for the Barcode library insert. A total of 20 standard ligation reactions were then set up as follows:

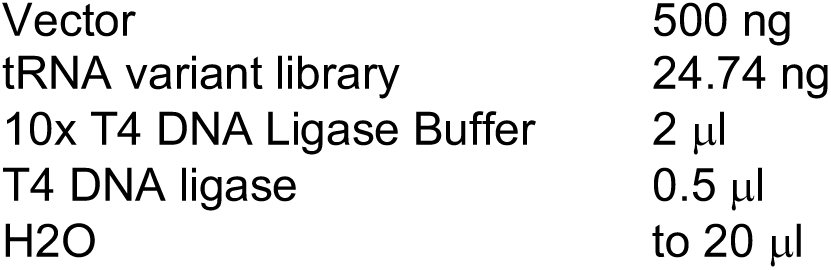

These reactions were incubated at 16°C for 16 hours. These were then pooled, co-purified, transformed, plated, harvested and final plasmids extracted by Qiagen plasmid Midiprep exactly as was done in the previous stage.

#### Barcode_library

TAACGAGGCGAAGACTAGTGCNNCANNGTNNAGNNNACNNAGCTCACGTCGGAGACGGACGTCTCCGCCCATGGACNNGANNNTCNNTCNNGANNGTAAATGTCTTCTGGCATTGC

Variable Bases, Buffer sequence, Dual BsmBI placeholder

^*^variable bases are 25% of each base at each site

#### tRNA variant library

CGTCTCtcgtcGGCTCGTNNGTCTNNNNNNNTGATTCTCGCTTAGGGTGCGAGAGGTCCCGGGNNNNNNNCCCGGACGAGCCCtGAGACG

tRNA^Pro^, Variable Bases

^*^variable bases are 82% of the wild-type base and 6% each of the three other possible bases at that location

### HEK-293T cell maintenance

HEK-293T cells (purchased from ATCC, ATCC-CRL-11268) were cultured in Dulbecco’s Modified Eagles Medium (DMEM) supplemented with 10% fetal bovine serum (FBS, E.U.-approved, South America origin, both from Gibco). Cells were maintained at 37°C 5% CO2 and passaged every 2-4 days at a ratio of 1:3-1:10. Cells tested negative for mycoplasma at least every 6 months using either a VenorGeM^®^ Mycoplasma Kit (Minerva Biolabs) according to manufacturer protocols, or using a set of primers from REF^25,26^ with Phusion^®^ High-Fidelity PCR Master Mix with GC Buffer (NEB). Cycling conditions for mycoplasma testing were:

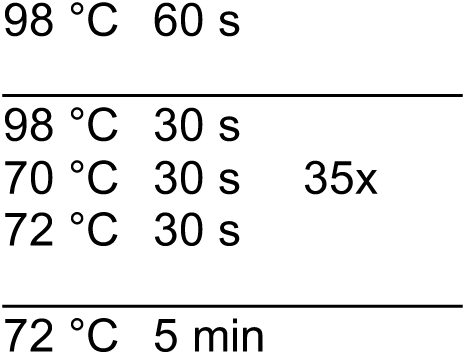

### Cell transfections and RNA extractions

For HEK-293T cell transfections cells were plated the night prior to transfection in 12-well tissue culture plates (Corning) in 500 μl DMEM + 10% FBS at numbers such that the next day they were at 60-80% confluence. Prior to transfection the media was removed and the wells washed once with 500 μl PBS. The media was then changed to 450 μl DMEM + 2% FBS. During this time, sufficient plasmid DNA such that at final volume (500 μl for these experiments) each plasmid would have a final concentration of 100 pM (50-500 ng each, depending on plasmid size) was brought up to 50 μl in Opti-MEM^TM^ (Gibco) containing 1.5 μg Polyethylenimine (PEI, Sigma). The solution was then mixed vigorously by vortexing for 10 seconds, let stand for 15 minutes at room temperature, and added drop-wise to the cells. Cells were then incubated for 4 hours at 37°C 5% CO_2_. Transfection media was then removed and replaced with fresh DMEM + 10% FBS and the cells left for an additional 20 hours (24 hours from the start of transfection). Wells were then harvested using 0.05% Trypsin-EDTA (Thermo Fisher Scientific). Following trypsinization, half of the total cells from each well were taken for flow cytometry. The remainder were spun down at 300g for 5 minutes, the supernatant removed, and the pellet snap frozen on dry ice. On thaw, RNA was extracted using ChargeSwitch^®^ Total RNA Cell Kit (Thermo Fisher Scientific) as per manufacturer instructions using an equal volume of 60°C elution buffer (E7) to input beads. For the tRNA screening library transfections, instead of ChargeSwitch^®^ extraction, 99% of cells were co-extracted for RNA and DNA using the AllPrep DNA/RNA/miRNA Universal Kit (Qiagen) as per manufacturer instructions. The remaining 1% was taken to check transfection efficiency by flow cytometry.

Experiments using the DtRNA^Gly^ backbones (Supplementary Fig. 7) were transfected as follows. DNA was prepared such that the final concentration of each plasmid would be at a final concentration of 156 pM (200-750 ng DNA). This was then brought up to 50 μl in OptiMEM with 6 μg PEI, vortexed to mix and incubated for 15 minutes at room temperature. The DNA-OptiMEM-PEI mix was then added to 250 000 HEK-293T cells in suspension in 450 μl DMEM + 5% FBS and plated into a well of a 24-well plate. Plates were then harvested 20-24 hours later for flow cytometry.

### RNA circularization assays

For all (+)CMV constructs, prior to circularization the 5’ cap was removed as this would otherwise inhibit the circularization reaction. To do so 12.5 U of RNA 5’ Pyrophosphohydrolase (RppH) was added to 2.5 μl of RNA in 1x ThermoPol^®^ Reaction Buffer (both from NEB) in a 25 μl total reaction volume. These were incubated for 1 hour at 37°C. RNA was then purified from the reactions using a ChargeSwitch^®^ Total RNA Cell Kit (Thermo Fisher Scientific) without lysis incubation or DNase treatment step and with a final elution volume of 12.5 μl. Both decapped (+)CMV and untreated (-)CMV samples were then circularized with reaction conditions as follows:

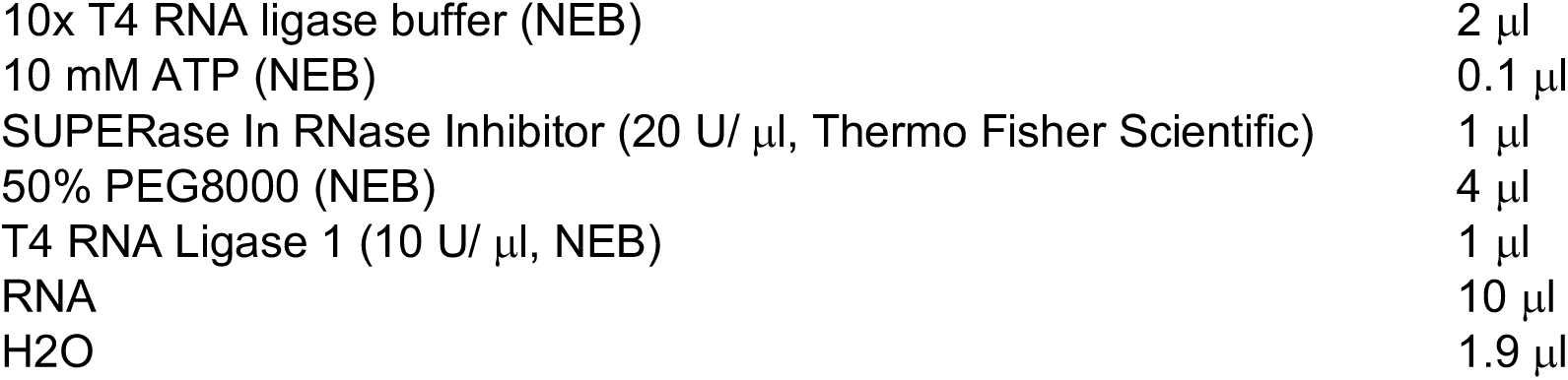

These reactions were incubated for 4 hours at room temperature then the RNA was re-purified using ChargeSwitch^®^ Total RNA Cell Kit as above with 12.5 μl elution volumes for the (+)CMV samples and 25 μl elution volumes for the (-)CMV. 6 μl of the resulting RNA was then reverse transcribed using the QuantiTect^®^ Reverse Transcription (RT) kit (Qiagen) as per manufacturer instructions with the specific primer ‘cRT-CTS2 nest_R’ (Supplemental Table 3) and with a 30 rather than a 15-minute incubation at 42°C. A nested PCR was then performed using Phusion^®^ High-Fidelity PCR Master Mix with GC Buffer (NEB) with 500 nM primer concentrations (‘cRT-sgRNA_nest_F’ + ‘cRT-CTS2_nest_R’ for 1^st^ PCR, cRT-sgRNA1_v2_F + ‘cRT-CTS2_R for 2^nd^ PCR, Supplemental Table 3) starting from 1 μl of the cDNA. 1 μl of a 1:10 dilution of the of the 1^st^ PCR was transferred into the 2nd PCR(20 μl total reaction volume). Cycling conditions were as follows:

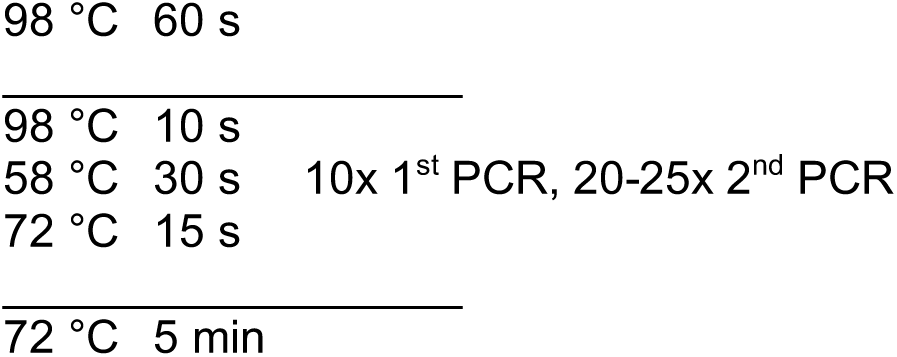

Following cycling, 2^nd^ PCR products were run on 2% agarose gels and nucleic acids visualized with GelRed^®^ (Biotium). Gel images were taken using a BioRad GelDoc^TM^ XR+ imager, with exposure times just below what would give saturated pixels (generally between 0.5 and 0.75 seconds). Image processing was done using GelAnalyzer2010a. First, automatic lane detection was performed, with lanes being manually adjusted in the case of errors. Next automatic peak identification was performed, and again manually curated. Then, rolling ball background subtraction (radius=25 pixels) was performed. Finally, the raw volume of the correctly processed peak was divided by the sum of the raw volume of all peaks to estimate the processing efficiency.

### Flow Cytometry

Flow cytometry was performed using either a BD LSRFortessa^TM^ cell analyzer or BD LSRII flow cytometer. ECFP was measured following 405 nm excitation with a 450/50 bandpass filter. EGFP was measured using a 488 nm excitation with a 530/30 (Fortessa) or 525/50 (LSRII) bandpass filter. iBlue was measured using a 640 nm excitation with a 670/14 bandpass filter. mCherry and mKate (for the intron release system) were measured with a 561 (Fortessa) of 532 nm (LSRII) excitation with a 610/20 bandpass filter.

### Deep sequencing library preparation - pDNA library preparation

The total pDNA fraction from the AllPrep DNA/RNA/miRNA Universal Kit (Qiagen) was first treated with 20 U Exonuclease V in NEBuffer 4 supplemented with 1 mM ATP at 37°C for 1 hour to remove genomic DNA. pDNA was then purified using Qiagen MinElute PCR purification columns with a 15 μl elution volume. Triplicate PCR amplifications were then performed for each sample as follows:

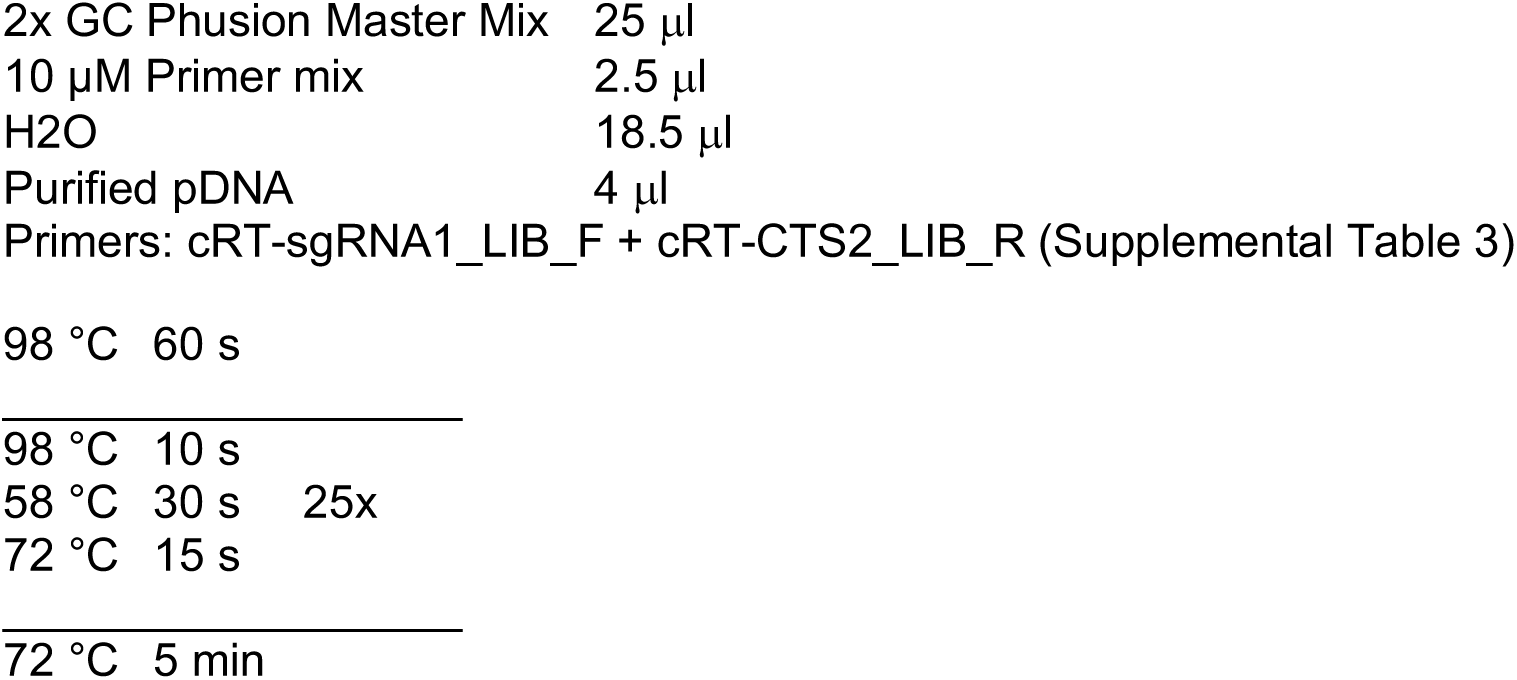

Triplicate amplicons were gel purified using a Qiagen gel extraction kit as per manufacturer instructions but with Qiagen MinElute PCR purification columns substituted for Qiaquick columns and a final elution volume of 15 μl buffer EB. Triplicate amplicons were then pooled at equimolar ratios for downstream steps.

### Deep sequencing library preparation - circRNA library preparation

First, the 5’ cap was removed from the (+)CMV libraries using RppH. For these reactions 37.5 U RppH was added to 750-1000 ng total RNA in 1x ThermoPol buffer (both from NEB) in a 75 μl reaction. These were incubated for 1 hour at 37°C. RNA was then purified from the reactions using ChargeSwitch^®^ Total RNA Cell Kit (Thermo Fisher Scientific) without the lysis incubation or DNase treatment step and with a final elution volume of 25 μl. Both the de-capped (+)CMV libraries and the untreated (-)CMV libraries were then circularized as follows:

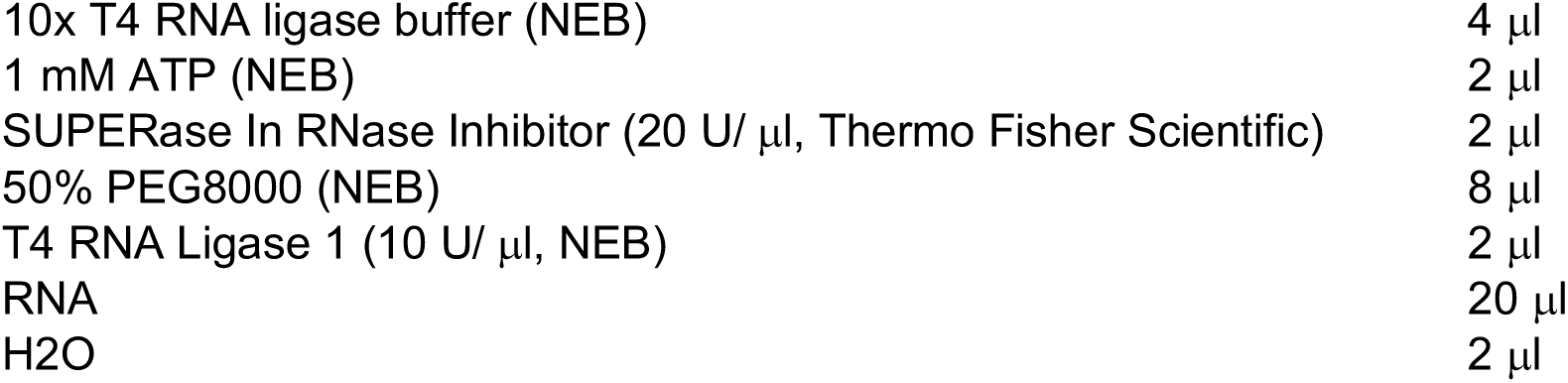

These reactions were incubated at room temperature for 4 hours at room temperature. RNA was then re-purified using ChargeSwitch^®^ Total RNA Cell Kit (Thermo Fisher Scientific) with a final elution volume of 25 μl. 1 μg of circularized RNA was then subjected to reverse transcription using the QuantiTect^®^ Reverse Transcription kit (Qiagen) as per manufacturer instructions with the specific primer ‘cRT-CTS2_nest_R’ (Supplemental Table 3) and with a 30 rather than a 15-minute incubation at 42°C. A first PCR amplification was then performed as follows:

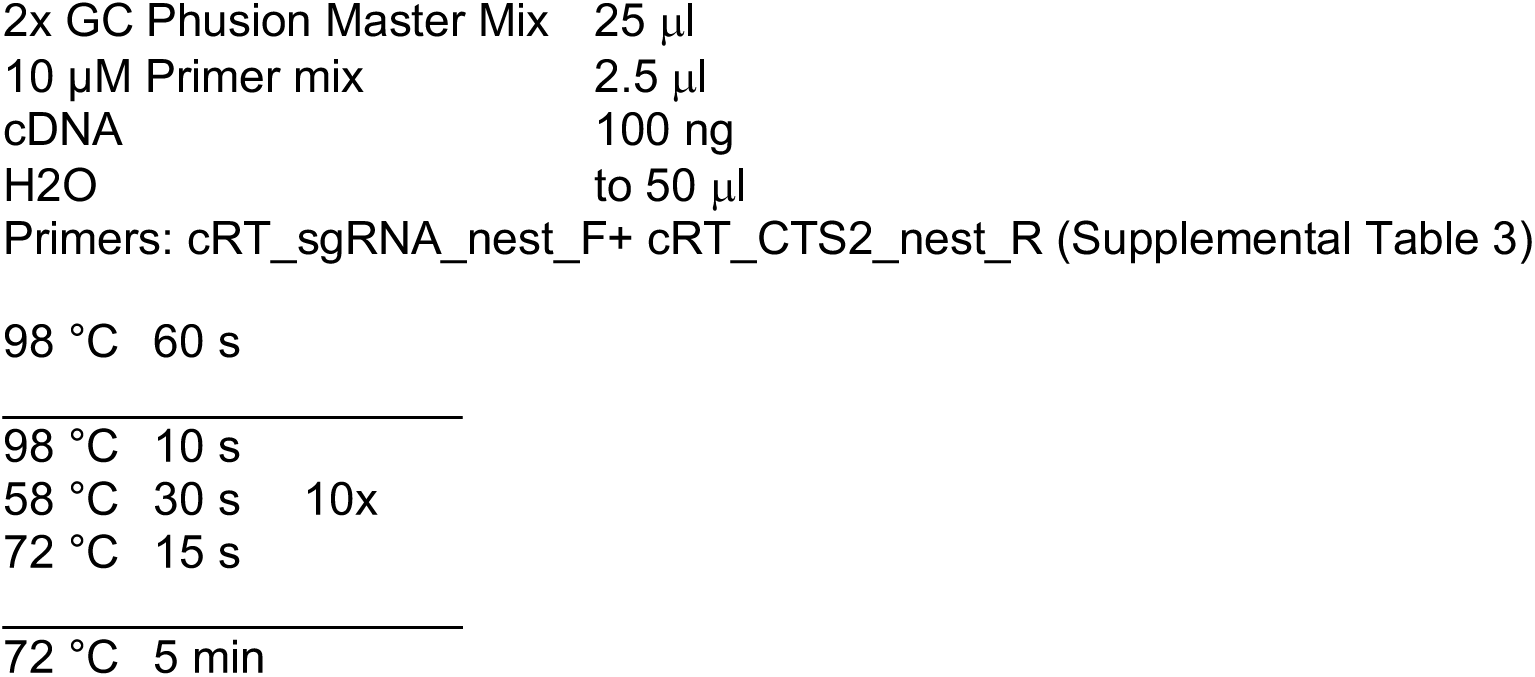

PCR products were diluted 1:10 in Ambion^®^ DEPC-treated water (Thermo Fisher Scientific) and used as template for a second PCR as follows:

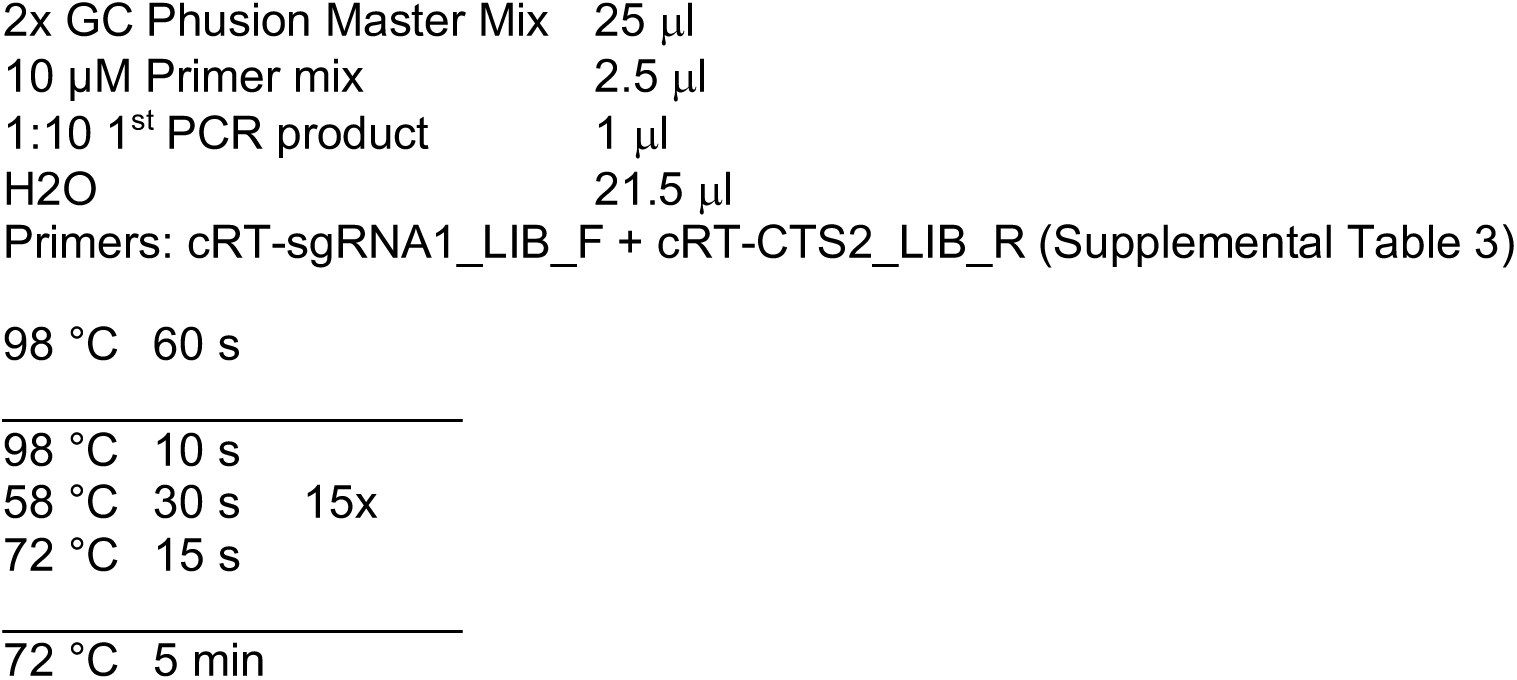

Following second PCR all samples were purified using Agencourt AMPure XP beads as per manufacturer instructions with 1.8x sample volumes of beads and a 25 μl final elution volume. Triplicate amplicons were then pooled at equimolar ratios for downstream processing.

### Deep sequencing library preparation - indexing and sequencing

Illumina indices were then added to both purified circRNA and pDNA libraries with another round of PCR amplification. For all samples the D508 was used as a forward index primer, while each sample was given a unique reverse index from D701-D712 (Supplemental Table 4). Reaction conditions were as follows:

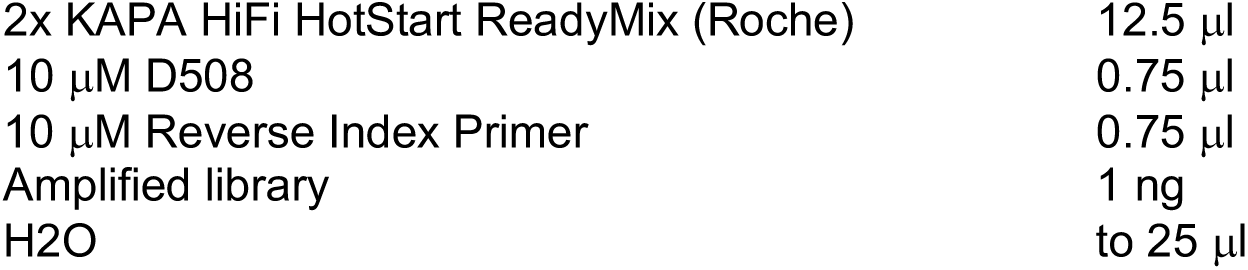

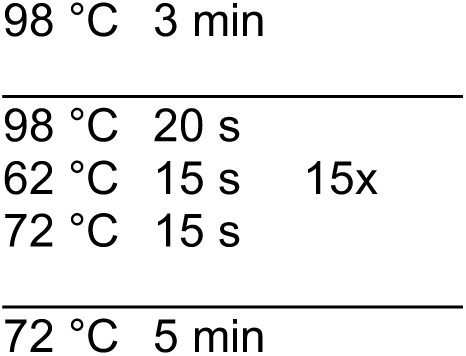

Following amplification, products were once again purified using Agencourt AMPure XP beads as per manufacturer instructions with 1.8x sample volumes of beads and a 25 μl final elution volume. Libraries were then quantified using a Qubit dsDNA HS (High Sensitivity) Assay Kit (Thermo Fisher Scientific) as per manufacturer instructions, and size distributions estimated by 2% agarose gel analysis using GelAnalyzer2010a. All libraries were then pooled and re-quantified for both concentration and size distribution and diluted to 4 nM. Finally, diluted libraries were sequenced using a MiSeq benchtop sequencer with a 300 cycle Reagent Kit v2 (Illumina) as per manufacturer protocols with 10% PhiX DNA spiked in.

### Quantitative RT-PCR

First, 3 μl RNA from each sample was treated with a TURBO DNA-free^TM^ Kit (Thermo Fisher Scientific) in a 6 μl total reaction as per manufacturer instructions to remove residual plasmid contamination. Next, RT was performed using the QuantiTect^®^ RT kit (Qiagen) with a 10 μl reaction volume using the supplied random primer with an incubation time of 30 rather than 15 minutes at 42°C. Quantitative PCR (qPCR) reactions were performed on the resulting cDNA using SsoAdvanced^TM^ Universal SYBR^®^ Green Supermix (Biorad) on a CFX384 Touch^TM^ Real-Time PCR Detection System (Biorad). All samples were measured with two primer pairs; ‘CTS2_qPCR-F’ with ‘sgRNA_common_qPCR-R’ for measuring sgRNA abundance, and ‘dCas9_qPCR-F’ with ‘dCas9_qPCR-R’ for dCas9-VP64 abundance as an internal normalization control (primer sequences available in Supplemental Table 3). Primers concentration was 250 nM and annealing temperature was 60°C.

### Data Analysis

All final analyses were performed in R (version 3.4.1). Statistical testing was performed using the ‘bayes.t.test’ function from the R library ‘BayesianFirstAid’ (version 0.1) with 30000 iterations for MCMC sampling. BEST tests^20^ were used as these are more information rich than classic *t*-tests and are robust to sample distribution and outliers. Tests were paired when relevant and unpaired in other cases (as specified for each test). For flow cytometric analysis a combination of functions from the R packages ‘flowCore’ (version 1.44.2) and custom scripts was used for basic processing (gating, plotting, summary statistics).

Analysis of deep sequencing data consisted of four stages; 1) pre-processing, 2) variant-barcode association, 3) barcode association refining and trimming, and 4) final analysis. First, the raw FASTQ files were quality trimmed using sickle (version 1.200) in paired-end mode with a quality threshold of 20 and a minimum read length of 70 bp. The (quality trimmed) paired-end reads were then merged into a single read based on sequence overlap using ‘<u>bbmerge-auto.sh</u>’ from BBMap (version 37.48) with k=60 on ‘strict’ mode.

As any read in the circRNA from a properly processed gRNA will not have the tRNA included in the read, an association map between the barcode sequences attached to the gRNAs and the sequence of the associated tRNA was necessary. To create the first iteration of such an association dictionary, we analysed the plasmid DNA sequencing results using a combination of ‘ShortRead’ (version 1.26.0), ‘Biostrings’ (version 2.36.4), and ‘plyr’ (version 1.8.4) together with custom scripts in R (version 3.2.1). To do so, we first identified reads in the expected amplicon size (between 185 and 200 bp). We next located the barcode sequences using ‘vcountPattern’ and ‘vmatchPattern’ allowing 1 mismatch (search patterns were “CNNCANNGTNNAGNNNACNN” for the 5’ barcode and “NNGANNNTCNNTCNNGANNGTAA” for the 3’ barcode). Barcode sequences were then retained if the potential barcode hit was <3 bp away from the expected location in the amplicon. Similarly, we identified tRNA sequences with the pattern “GGCTCGTNNGTCTNNNNNNNTGATTCTCGCTTAGGGTGCGAGAGGTCCCGGGNNNNNNNCCCGGACGAGCCC” now allowing 3 mismatches.

Reads which had identifiable barcodes (5’ and 3’) and a tRNA sequence of correct length and which did not have any ambiguous bases were retained. Next, reads with identical barcode and tRNA sequence were collapsed to obtain read counts. Finally, for any barcode pair with inconsistent tRNA sequences, inconsistent sequences with 2 bp or less difference from the most abundant read and less than 50% of the most abundant read were assumed to be sequencing errors and merged into the most abundant reads count. If more differences were present, the other differing sequences were ignored if they had a read abundance of 10% or less of the most abundant read. If inconsistent reads had more than 2 bp difference from the most abundant read and were >10% the abundance of the most abundant read, all reads of this barcode pair were removed from the dictionary.

Frequencies for each mutation in the overall variant library in each experiment was calculated as the number of reads for that tRNA variant in the retained reads divided by the total number of retained reads. Expected frequency (f) for 0 to 10 mutations (n) was calculated as _16_C_n_(f^(16-n)^(1-f)^n^. The mean squared error (m.s.e.) was calculated between the observed frequencies of each number of mutations and the predicted in increments of 0.033% across the probable range of values and the best fit chosen which minimized m.s.e. The distribution of mutations across nucleotides was calculated as the number of reads with a mutation at a given site divided by the total number of reads.

For the circRNA, all merged reads with a total length of 50 bp or more were considered. We next identified the tRNA, 5’ and 3’ barcodes as was done for the pDNA for the (-)CMV libraries. For (+)CMV libraries we required only the first 9 variable bases of the 5’ barcode to be present (a pattern of “CGGTGCNNCANNGTNNAGNNN” for the 5’ barcode) as the barcode was partially truncated in a majority of the sequences, possibly due to degradation during the 5’ cap removal. In the case of the truncated requirement, pDNA barcode associations were also collapsed to retain only those with unique truncated barcode to tRNA sequence associations. Following identification of barcodes and tRNA sequences, we next classified each read based on its degree of processing. To be classified as processed, a read had to have a total length less than 150 bp, have one but not the other barcode sequence (ie either 5’ or 3’), and not have an identifiable tRNA sequence. Reads were classified as fully unprocessed if both barcodes and tRNA sequence were present and the overall length was 150 bp or greater. Finally, they were classified as partially processed if total length was 150 bp or greater, one barcode was present and a tRNA sequence.

We next created barcode-tRNA sequence dictionaries based on the unprocessed circRNA as was done for the pDNA. In this case dictionaries were separate for the 5’ and 3’ barcodes as in the case of partial processing both barcodes were not necessarily present in the same read. To obtain our final barcode-tRNA dictionary, we then compared the circRNA dictionaries to the pDNA dictionaries. In cases of disagreement these were resolved as previously done within the pDNA or circRNA (i.e. merging those which were very similar, ignoring very low abundance disagreeing reads, and removing barcodes with ambiguous associations). Finally, we only retained those associations which had been observed in at least 3 reads and had at least one processed and one unprocessed read in the circRNA dataset.

Processing efficiency was calculated for each tRNA variant as 100^*^(processed reads/all reads). To determine the processing ability of each mutation a binomial distribution was inferred for each replicate based on the number of processed reads and the number of total reads using the R function ‘dbinom’ in the range of 1-100% with 1% increments. These distributions were averaged across the three replicates to get an overall probability distribution. The maximum likelihood of the combined binomial distributions was used as the estimated processing efficiency. Significance values were calculated by determining the area of overlap of the two probability distributions to be compared. The effect of each mutation on promoter activity was calculated as log_2_(cRNA_freq_/pDNA_freq_). Statistical testing for differences in promoter activity were calculated using paired BEST tests comparing each mutation to wild-type.

## ACKNOWLEDGEMENTS

D.J.H.F.K. is funded by a CIHR Postdoctoral Fellowship. T.A.F. was supported by MRC (G0902418), BBSRC (BB/N006550/1) and Wellcome Trust ISSF (105605/Z/14/Z). Y.S.M. is funded by the WIMM prize studentship, Clarendon Scholarship and Christopher Welch Scholarship. Q.R.V.F. was funded by a Wellcome Trust PhD Studentship. M.J. acknowledges funding from the University of Oxford and the EPSRC & BBSRC Centre for Doctoral Training in Synthetic Biology (grant EP/L016494/1). T.A.M is funded by Medical Research Council (MRC, UK) Molecular Haematology Unit Grant MC_UU_12009/6.

## AUTHOR CONTRIBUTIONS

D.J.H.F.K. and T.A.F. conceived the project, designed the experiments, interpreted the results and wrote the manuscript. D.J.H.F.K. performed all experiments. Y.S.M. helped with the tRNA screen design and analysis framework. M.J. assisted with cloning and qPCR optimization. Q.R.V.F. assisted with the design of the reporter assays and plasmid preparation. H.B. cloned wild-type tRNA^Gln^ constructs. T.A.M. provided project guidance. All authors provided input on the manuscript.

## REFERENCES

1. Cong, L. et al. Multiplex genome engineering using CRISPR/Cas systems. Science 339, 819–823 (2013).

2. Mali, P. et al. RNA-guided human genome engineering via Cas9. Science 339, 823–826 (2013).

3. White, R. J. Transcription by RNA polymerase III: more complex than we thought. Nat. Rev. Genet. 12, 459–463 (2011).

4. Nissim, L., Perli, S. D., Fridkin, A., Perez-Pinera, P. & Lu, T. K. Multiplexed and programmable regulation of gene networks with an integrated RNA and CRISPR/Cas toolkit in human cells. Mol. Cell 54, 698–710 (2014).

5. Shechner, D. M., Hacisuleyman, E., Younger, S. T. & Rinn, J. L. Multiplexable, locus-specific targeting of long RNAs with CRISPR-Display. Nat. Methods 12, 664–670 (2015).

6. Kiani, S. et al. CRISPR transcriptional repression devices and layered circuits in mammalian cells. Nat. Methods 11, 723–726 (2014).

7. Xu, L., Zhao, L., Gao, Y., Xu, J. & Han, R. Empower multiplex cell and tissue-specific CRISPR-mediated gene manipulation with self-cleaving ribozymes and tRNA. Nucleic Acids Res. (2016). doi:10.1093/nar/gkw1048

8. Lodish, H. et al. Molecular Cell Biology. (W. H. Freeman, 2000).

9. Xie, K., Minkenberg, B. & Yang, Y. Boosting CRISPR/Cas9 multiplex editing capability with the endogenous tRNA-processing system. Proc. Natl. Acad. Sci. U. S. A. 112, 3570–3575 (2015).

10. Qi, W. et al. High-efficiency CRISPR/Cas9 multiplex gene editing using the glycine tRNA-processing system-based strategy in maize. BMC Biotechnol. 16, 58 (2016).

11. Mefferd, A. L., Kornepati, A. V. R., Bogerd, H. P., Kennedy, E. M. & Cullen, B. R. Expression of CRISPR/Cas single guide RNAs using small tRNA promoters. RNA N. Y. N 21, 1683–1689 (2015).

12. Wei, Y. et al. CRISPR/Cas9 with single guide RNA expression driven by small tRNA promoters showed reduced editing efficiency compared to a U6 promoter. RNA N. Y. N 23, 1–5 (2017).

13. Port, F. & Bullock, S. L. Augmenting CRISPR applications in Drosophila with tRNA-flanked sgRNAs. Nat. Methods (2016). doi:10.1038/nmeth.3972

14. Sharp, S., DeFranco, D., Dingermann, T., Farrell, P. & Söll, D. Internal control regions for transcription of eukaryotic tRNA genes. Proc. Natl. Acad. Sci. U. S. A. 78, 6657–6661 (1981).

15. Levinger, L. et al. Matrices of paired substitutions show the effects of tRNA D/T loop sequence on Drosophila RNase P and 3’-tRNase processing. J. Biol. Chem. 273, 1015–1025 (1998).

16. Li, C., Qian, W., Maclean, C. J. & Zhang, J. The fitness landscape of a tRNA gene. Science 352, 837–840 (2016).

17. Hull, M. W., Erickson, J., Johnston, M. & Engelke, D. R. tRNA genes as transcriptional repressor elements. Mol. Cell. Biol. 14, 1266–1277 (1994).

18. Kendall, A. et al. A CBF5 mutation that disrupts nucleolar localization of early tRNA biosynthesis in yeast also suppresses tRNA gene-mediated transcriptional silencing. Proc. Natl. Acad. Sci. 97, 13108–13113 (2000).

19. Haurwitz, R. E., Jinek, M., Wiedenheft, B., Zhou, K. & Doudna, J. A. Sequence- and structure-specific RNA processing by a CRISPR endonuclease. Science 329, 1355–1358 (2010).

20. Kruschke, J. K. Bayesian estimation supersedes the t test. J. Exp. Psychol. Gen. 142, 573–603 (2013).

21. BååthR, R. Bayesian First Aid: A Package that Implements Bayesian Alternatives to the Classical ^*^.test Functions in R. in UseR! 2014 - the International R User Conference (2014).

22. Ferry, Q. R. V., Lyutova, R. & Fulga, T. A. Rational design of inducible CRISPR guide RNAs for de novo assembly of transcriptional programs. Nat. Commun. 8, 14633 (2017).

23. Knapp, D. J. H. F. et al. Factors influencing the sensitivity and specificity of conventional sequencing in human immunodeficiency virus type 1 tropism testing. J. Clin. Microbiol. 51, 444–451 (2013).

24. Knapp, D. J. H. F. et al. ‘Deep’ sequencing accuracy and reproducibility using Roche/454 technology for inferring co-receptor usage in HIV-1. PloS One 9, e99508 (2014).

25. Uphoff, C. C. & Drexler, H. G. Comparative PCR analysis for detection of mycoplasma infections in continuous cell lines. In Vitro Cell. Dev. Biol. Anim. 38, 79–85 (2002).

26. Uphoff, C. C. & Drexler, H. G. Detecting Mycoplasma contamination in cell cultures by polymerase chain reaction. Methods Mol. Med. 88, 319–326 (2004).

